# Comparative analysis of bile-salt degradation in *Sphingobium* sp. strain Chol11 and *P. stutzeri* Chol1 reveals functional diversity of β-proteobacterial steroid degradation enzymes and suggests a novel reaction sequence for side-chain degradation involving a hydroxylation step

**DOI:** 10.1101/2021.06.24.449856

**Authors:** Franziska Maria Feller, Phil Richtsmeier, Maximilian Wege, Bodo Philipp

## Abstract

The reaction sequence for aerobic degradation of bile salts by environmental bacteria resembles degradation of other steroid compounds. Recent findings show that bacteria belonging to the *Sphingomonadaceae* use a pathway variant for bile-salt degradation. This study addresses this so-called Δ^4,6^ -variant by comparative analysis of unknown degradation steps in *Sphingobium* sp. strain Chol11 with known reactions found in *Pseudomonas stutzeri* Chol1. Investigations with strain Chol11 revealed an essential function of the acyl-CoA dehydrogenase Scd4AB for growth with bile salts. Growth of the *scd4AB* deletion mutant was restored with a metabolite containing a double bond within the side chain which was produced by the Δ^22^-acyl-CoA dehydrogenase Scd1AB from *P. stutzeri* Chol1. Expression of *scd1AB* in the *scd4AB* deletion mutant fully restored growth with bile salts while expression of *scd4AB* only enabled constricted growth in *P. stutzeri* Chol1 *scd1A* or *scd1B* deletion mutants. Strain Chol11 Δ*scd4A* accumulated hydroxylated steroid metabolites which were degraded and activated with coenzyme A by the wild type. Activities of five Rieske type monooxygenases of strain Chol11 were screened by heterologous expression and compared to the B-ring cleaving KshAB_Chol1_ from *P. stutzeri* Chol1. Three of the Chol11 enzymes catalyzed B-ring cleavage of only Δ^4,6^ -steroids while KshAB_Chol1_ was more versatile. Expression of a fourth KshA homolog, Nov2c228 led to production of metabolites with hydroxylations at an unknown position. These results indicate functional diversity of β-proteobacterial enzymes for bile-salt degradation and suggest a novel side-chain degradation pathway involving an essential ACAD reaction and a steroid hydroxylation step.

**Importance:** This study highlights the biochemical diversity of bacterial degradation of steroid compounds in different aspects. First, it further elucidates an unexplored variant in the degradation of bile-salt side chains by Sphingomonads, a group of environmental bacteria that is well-known for their broad metabolic capabilities. Moreover, it adds a so-far unknown hydroxylation of steroids to the reactions Rieske monooxygenases can catalyze with steroids. Additionally, it analyzes the first proteobacterial ketosteroid-9α-hydroxylase and shows that this enzyme is able to catalyze side reactions with non-native substrates.

## Introduction

While steroid biosynthesis is apparently restricted to very few prokaryotic phyla (1, 2), steroids from eukaryotic organisms can serve as energy and carbon source for many different environmental bacteria (3, 4). Steroids have a nucleus of three C_6_- and one C_5_-ring, named A-D, and differ in the number and position of functional groups such as hydroxy groups and the presence of a side chain that may be attached to the C_5_ ring D. Bile salts comprise steroidal compounds involved in digestion and signaling of e.g. vertebrates and are excreted into environment in large amounts (5). Mammalian bile salts carry a carboxylic side chain as well as one to three hydroxy groups (e.g. cholate, I in Fig 1) attached to the steroid nucleus. Bacterial steroid degradation is modular and can be divided in four phases (3, 4, 6, 7): 1) Oxidation of the A-ring (Fig 1A), 2) side-chain degradation (Fig 1B), 3) degradation of rings A and B by oxygenation (aerobic) (Fig 1C) or hydrolysis (anaerobic; not shown (8, 9)), and 4) hydrolytic degradation of the remaining rings C and D. While the last module is very conserved (4, 10), the other modules can differ depending on the group of bacteria. In particular, the presence of a 7-OH in some bile salts enables a only scarcely elucidated variation, called Δ^4,6^-variant, of initial oxidation reactions (phase 1), that is found in many *Sphingomonadaceae*, which are also proposed to employ a so-far unknown side-chain degradation mechanism (11–13 (preprint)).

**Fig 1.**
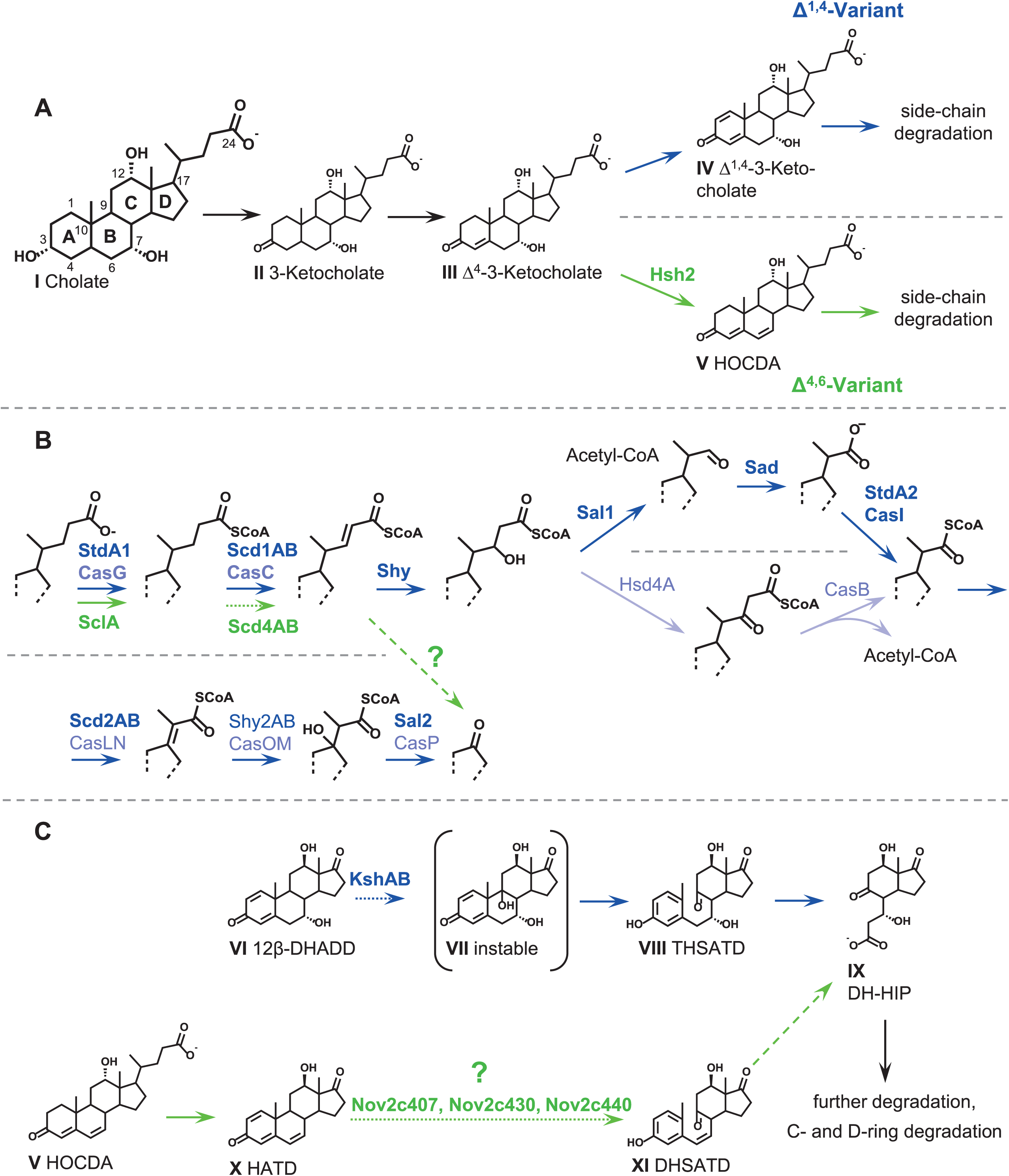
Overview over the degradation of cholate by bacteria. (A) A-ring oxidation and diversion into the Δ^1,4^- and Δ^4,6^-variant by either introduction of a second double bond into the A-ring (Δ^1,4^-variant, blue) or the elimination of water at C7 by key enzyme 7α-hydroxysteroid dehydratase (Δ^4,6^-variant, green). (B) Side-chain degradation by *P. stutzeri* Chol1 (dark blue) or *R. jostii* RHA1 (light blue) and known side-chain degradation steps of *Sphingobium* sp. strain Chol11 (green). (C) Section of the degradation of the steroid nucleus via the 9,10-*seco* pathway as found in *P. stutzeri* Chol1 (blue) and potential channelling of Δ^4,6^-intermediates to common C- and D-ring degradation (green). Dotted lines, elucidated in this study; broken lines, suggested reactions; bold names, experimentally verified; light names, bioinformatically predicted. Abbreviations: V, HOCDA (12α-hydroxy-3-oxo-4,6-choldienoic acid); VI, 12β-DHADD (7α,12β-dihydroxy-androsta-1,4-diene-3,17-dione); VII, 7α,9α,12β-trihydroxy-androsta-1,4-diene-3,17-dione (unstable); VIII, THSATD (3,7,12-trihydroxy-9,10-*seco*-androsta-1,3,5(10)-triene-9,17-dione); IX, DH-HIP (3’,7-dihydroxy-H-methyl-hexahydro-indanone-propanoate); X, HATD (12-hydroxy-androsta-1,4,6-triene-3,17-dione); XI, DHSATD (3,12β-dihydroxy-9,10-*seco*-androsta-1,3,5(10),6-tetraene-9,17-dione).

In most model strains including *Rhodococcus jostii* RHA1, *Comamonas testosteroni,* and *Pseudomonas stutzeri* Chol1, aerobic bile-salt degradation proceeds via the so-called 9,10-*seco* pathway involving intermediates with a Δ^1,4^-3-keto structure of the steroid skeleton, which are formed by oxidation of the steroidal A-ring (blue in Fig 1 A) (3, 14, 15). In this first phase, the hydroxy group at C3 is oxidized to a keto group and two double bonds at C1 and C4 are introduced into the A-ring, which leads to Δ^1,4^-3-ketocholate (IV in Fig 1) for the bile salt cholate (I in Fig 1) (16). In second phase, degradation of the C_5_ carboxylic side chain proceeds similar to the β-oxidation of fatty acids, and acetyl-CoA and propionyl-CoA are released consecutively (Fig 1B) (16–18). The side chain is first activated with CoA by a CoA-ligase such as StdA1 in *Pseudomonas putida* DOC21 (19), then a double bond is introduced between C22 and C23 by an acyl-CoA-dehydrogenase (ACAD) such as Scd1AB in *P. stutzeri* Chol1 (20). To this double bond, water is added by a hydratase such as Shy1 to gain a β-hydroxy group (17). While acetyl-CoA is released by β-oxidation in actinobacteria such as *R. jostii* RHA1 (14), aldolytic cleavage similar to β-oxidation can be found in proteobacteria such as *P. stutzeri* Chol1 (17, 21). For this, acetyl-CoA is released by the aldolase Sal1. This results in a shortened side chain with an aldehyde group, which is then oxidized to a carboxyl group by aldehyde dehydrogenase Sad (17). The resulting intermediates such as 7,12-dihydroxy-3-oxo-pregna-1,4-diene-carboxylate (DHOPDC) (XII in Fig 2) have a C_3_ carboxylic side chain (16, 21, 22), which is then released as propionyl-CoA. For this, another cycle of aldolytic cleavage in actinobacteria as well as proteobacteria is catalyzed by consecutive CoA-activation by StdA2 (19), dehydrogenation by the second ACAD Scd2AB (20, 22), hydroxylation and aldolytic cleavage resulting in C_19_-steroids called androsta-1,4-diene-3,17-diones (ADD) such as 7,12β-dihydroxy-ADD (12β-DHADD, VI) for cholate (I) (14, 18, 23). In the third phase of degradation, rings A and B are cleaved, which occurs simultaneously or consecutively to side-chain degradation in actinobacteria (24) or proteobacteria, respectively (22). First, the B-ring is cleaved by monooxygenase KshAB, which introduces a hydroxy group at C9 (Fig 1C). The resulting 9α-hydroxy-ADD (VII) spontaneously reacts to the name-giving 9,10-*seco* intermediates such as 3,7,12-trihydroxy-9,10-*seco*-androsta-1,3,5-triene-9,17-dione (THSATD, VIII) driven by the aromatization of the A-ring. Further degradation is achieved by oxygenation and *meta*-cleavage of the A-ring and hydrolytic cleavage of the A-ring residue resulting intermediates consisting of former rings C and D called *H*-methyl-hexahydro-indanone-propanoates (HIP) such as 3’,7-dihydroxy-HIP (DH-HIP, IX) for cholate (blue in Fig 1C) (3, 4). At this stage of degradation, intermediates from differently hydroxylated bile salts are channeled to one common intermediate in *P. stutzeri* Chol1 (20). Further degradation of HIPs proceeds via β-oxidation and hydrolytic cleavages resulting in acetyl-CoA, succinyl-CoA, and propionyl-CoA (25, 26).

**Fig 2.**
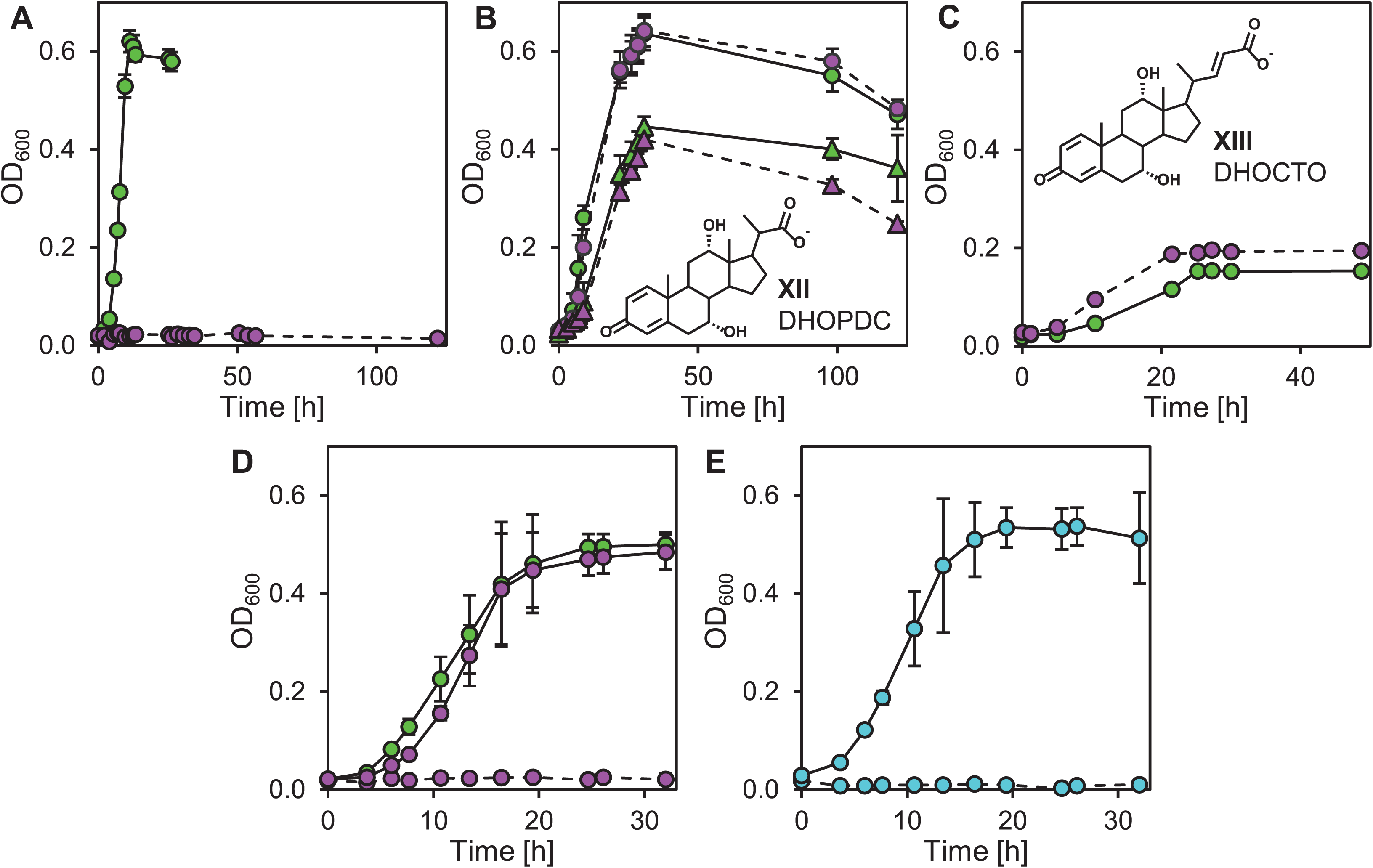
Phenotype of *Sphingobium* sp. strain Chol11 Δ*scd4A*. **(A)** Growth of strain Chol11 Δ*scd4A* (dotted lines, red) and wt (solid lines, green) with 1 mM cholate. **(B)** Growth of strain Chol11 Δ*scd4A* (dotted lines, red) and wt (solid lines, green) with steroids with shortened C_3_-side chain (XII, 7,12-dihydroxy-3-oxo-pregna-1,4-diene-carboxylate, DHOPDC; 1 mM, triangles) or no side chain (12β-DHADD, VI in Fig 1; 2 mM, circles). **(C)** Growth of strain Chol11 Δ*scd4A* (dotted lines, red) and wt (solid lines, green) with 7,12-dihydroxy-3-oxo-chol-1,4,22-triene-oate (DHOCTO, XIII; about 0.2 mM). **(D)** Complementation of strain Chol11 Δ*scd4A*. Growth of strain Chol11 Δ*scd4A* pBBR1MCS-5::*scd4A* (solid lines, red), Δ*scd4A* pBBR1MCS-5 (dotted lines, red), and wt pBBR1MCS-5::*scd4AB* (solid lines, green) with 1 mM cholate. **(E)** Heterologous complementation of strain Chol11 Δ*scd4A* with *scd1AB* of *P. stutzeri* Chol1. Growth of strain Chol11 Δ*scd4A* pBBR1MCS-5::*scd1AB_Chol1_* (solid lines, turquois) and Δ*scd4A* pBBR1MCS-5::*scd1A_Chol1_* (dotted lines, turquois) with 1 mM cholate. Error bars indicate standard deviations and may not be visible if too small (n=3).

In contrast to this well elucidated pathway, some steps of 7-hydroxy bile-salt degradation proceed differently in *Sphingobium* sp. strain Chol11, which uses the aforementioned Δ^4,6^-variant. While the first steps of A-ring oxidation also lead to Δ^4^-3-keto intermediates such as Δ^4^-3-ketocholate (III in Fig 1) (11, 27), in the next step the hydroxy group at C7 is eliminated by 7α-hydroxy steroid dehydratase Hsh2 (green in Fig 1A) (28). This leads to Δ^4,6^-3-keto intermediates such as 12-hydroxy-3-oxo-chol-4,6-dienoate (HOCDA, V) with a double bond in the B-ring. This pathway variation has also been found in other *Sphingomonadaceae* such as *Novosphingobium tardaugens* NBRC16725 and *Novosphingobium aromaticivorans* F199 (13 (preprint)). The transient accumulation of Δ^4,6^-derivatives of ADDs such as 12-hydroxy-androsta-1,4,6-triene-3,17-dione (HATD, X) in the supernatants of cultures of strain Chol11 (11) indicates that side-chain degradation is the next step of degradation. The side chain is CoA-activated by CoA-ligase SclA in strain Chol11 and in enzyme assays introduction of a double bond into the side chain was observed (12). However, the position of this double bond as well as further side-chain degradation has not been elucidated yet, and strain Chol11 does apparently not harbor homologs of all proteins involved in either cycle of side-chain cleavage of *P. stutzeri* Chol1 or *R. jostii* RHA1 (12). Differential proteome analyses of strain Chol11 adapted to growth with cholate versus ADDs without side chain together with bioinformatical analyses of several bile-salt degrading *Sphingomonadaceae* led to the identification of a potential side-chain degradation cluster (Fig S1) (13 (preprint)). The core set of genes in this cluster consists of SclA (12), two adjacent putative ACADs, a Rieske monooxygenase with similarity to KshA and an amidase. In strain Chol11, the cluster additionally encodes an amidase that is able to cleave conjugated bile salts (13 (preprint)), two adjacent thioesterase family proteins, putative steroid dehydrogenases as well as transporters. Further degradation of the steroid skeleton is also unknown, but proteomic and bioinformatical analyses strongly indicated that the degradation of the steroid nucleus also proceeds via the 9,10-*seco* pathway (12, 13 (preprint)). However, no *seco-* steroids with Δ^4,6^-structure could so far be detected in culture supernatants of strain Chol11 growing with cholate.

The goal of this study was to elucidate bile-salt side-chain degradation in *Sphingobium* sp. strain Chol11 by a functional analysis of the gene cluster predicted in the proteomic study. For this, deletion mutants of several genes in the putative side-chain degradation gene cluster were constructed. Additionally, comparative analyses with the model organism *P. stutzeri* Chol1, in which bile-salt degradation is very well elucidated, were performed e.g. by heterologous expression of candidate genes in suitable mutants of *P. stutzeri* Chol1 (17, 21).

## Results

### Nov2c221/222 is involved in steroid C_5_ side-chain degradation

The next plausible step of side-chain degradation after CoA-activation would be the introduction of a double bond at C22 by an ACAD, which has been observed for HOCDA-CoA in enzyme assays (12). In the side-chain degradation cluster of *Sphingobium* sp. strain Chol11, two subunits for an ACAD (Nov2c221 and Nov2c222) are encoded by adjacent genes. This gene synteny is also known for other ACADs involved in steroid metabolism, that are heterotetramers of two ACAD subunits (29). In a reciprocal BLASTp analysis, Nov2c221 and Nov2c222 were indeed annotated as the two subunits of the HIP-ACAD, Scd3A (20) (also called ScdD1 (26)) and Scd3B (ScdD2), with 58 % and 40 % identity to the homologs of *P. stutzeri* Chol1, respectively (12). This is also reflected in a phylogenetic tree of steroid-degradation ACADs (Fig S2), in which Nov2c221 and Nov2c222 cluster with the α- and β-subunits, respectively, of HIP-ACADs. However, due to their location and higher abundance in cholate-versus ADD-grown cells (13 (preprint)), a role in side-chain degradation seems to be more likely, although the similarities to the respective subunits of C_5_- and C_3_-side chain ACADs Scd1AB and Scd2AB were much lower with only up to 33 % (Table 1).

**Table 1.**
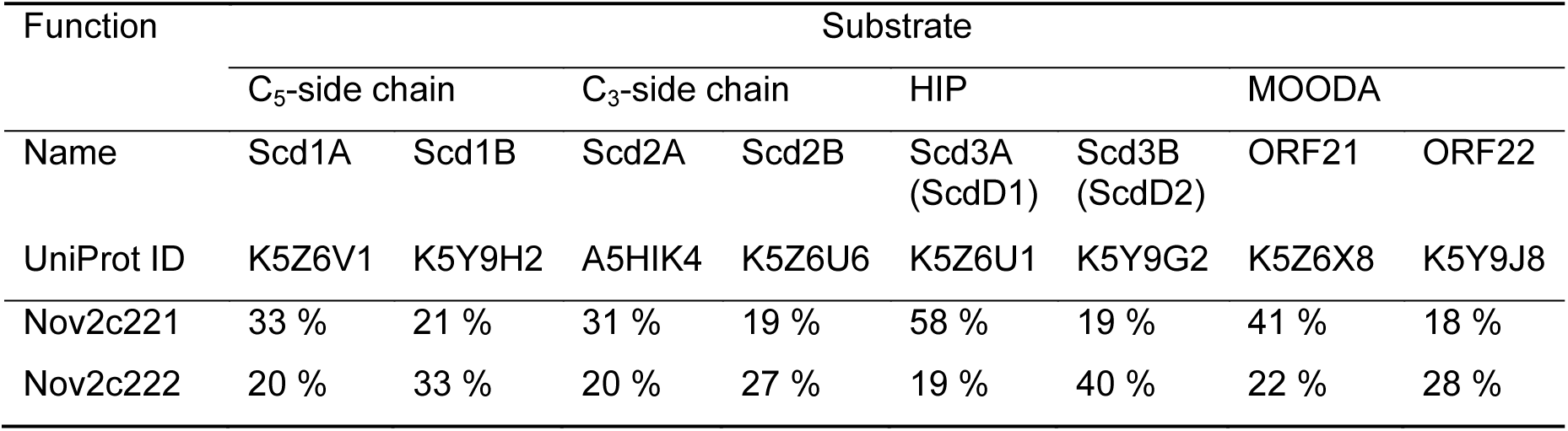
Identities of Nov2c221 and Noc2c222 of *Sphingobium* sp. strain Chol11 to the subunits of different steroid-degradation ACADs of *P. stutzeri* Chol1. Names in *C. testosteroni* CNB-2 are given in parentheses if different. MOODA, 4-methyl-5-oxo-octanedioate; linear intermediate in the degradation of the C- and D-ring.

To test the function of Nov2c221 and Nov2c222, an unmarked deletion mutant lacking the gene for the presumptive catalytically active α-subunit, Nov2c221, was constructed. The deletion mutant strain Chol11 Δ*nov2c221* did not grow with cholate as the only carbon source (Fig 2A).

Growth with cholate could be restored by expression of a plasmid borne copy of *nov2c221* (Fig 2D) excluding possible downstream effects especially in this operon-like structure. The mutant strain could grow with cholate degradation intermediates with a shortened or without side chain, namely DHOPDC (XII in Fig 2) and 12β-DHADD (VI in Fig 1) (Fig 2B). With glucose as the only carbon source, strain Chol11 Δ*nov2c221* grew very similarly to the wild type (not shown). Cholate transformation experiments with dense cell suspensions showed that the deletion mutant depleted cholate with a strongly decreased rate compared to the wild type (Fig S3A, complete depletion of cholate within 30 h instead of 4 h). The deletion mutant constantly formed HOCDA (V in Fig 1, Fig S3B) within these 30 h, that reached 100x elevated concentrations compared to the maximum concentrations that were transiently accumulated by the wild type. These results showed that the deletion mutant harbored functional A-ring oxidation and water elimination from the B-ring as well as the degradation of the steroid ring including a shortened side chain but was unable to degrade the C_5_-side chain.

In the next step, we tried to restore the growth of strain Chol11 Δ*nov2c221* by providing a substrate that already has a double bond in the side chain at C22, namely 7,12-dihydroxy-3-oxo-chol-1,4,22-triene-oate (DHOCTO, XIII in Fig 2). Both the wild type and the deletion mutant were able to grow with and degrade DHOCTO (Fig 2C).

### Nov2c221/222 and C_5_ side-chain ACAD Scd1AB can replace each other in cross-complementation experiments with different efficiencies

As DHOCTO is formed by the ACAD Scd1AB in *P. stutzeri* Chol1, we tested whether cross-complementation of the individual enzymes of strains Chol1 and Chol11 was possible. For this, deletion mutants of *P. stutzeri* Chol1 lacking either subunit (20) were heterologously complemented with *nov2c221* and *nov2c222. P. stutzeri* Chol1 strains Δ*scd1A* and Δ*scd1B* carrying plasmid-borne copies of *nov2c221* and *nov2c222* in combination were able to grow with cholate as the only carbon source (Fig 3A) while the vector control was not. The lag phase of these cross-complemented strains was strongly, but inconsistently increased for these strains: lag phases of *P. stutzeri* Chol1 Δ*scd1A* pBBR1MCS-5::*nov2c221-222* were varying from 24 h to up to 250 h and the lag phases of *P. stutzeri* Chol1 Δ*scd1B* pBBR1MCS- 5::*nov2c221-222* always were about 250 h (Fig 3A shows representative examples). In addition, growth rates were reduced and final OD_600_ values were diminished to 0.4 instead of 0.7 in comparison to the wild type (23). In the culture supernatant of the cross-complemented strains DHOPDC (XII in Fig 2) with shortened C_3_-side chain as well as THSATD (VIII in Fig 1) without side chain were detected as degradation intermediates (shown for *P. stutzeri* Cho1 Δ*scd1A* pBBR1MCS-5::*nov2c221-222* in Fig 3B). This indicates functional two-step side-chain degradation in the cross-complemented strains. Additionally, *nov2c221* proposedly encoding the α-subunit was expressed in *P. stutzeri* Chol1 Δ*scd1A* but was not able to restore growth on its own (data not shown). The other way around, *scd1AB_Chol1_* were expressed in strain Chol11 Δ*nov2c221*. The respective strain carrying the plasmid pBBR1MCS-5::*scd1AB_Chol1_* grew very similarly to the wild type and, thus, without detectable lag phase with cholate as the only carbon source (Fig 2E). In contrast to this, *scd1A_Chol1_* alone was not able to restore growth (Fig 2E).

**Fig 3.**
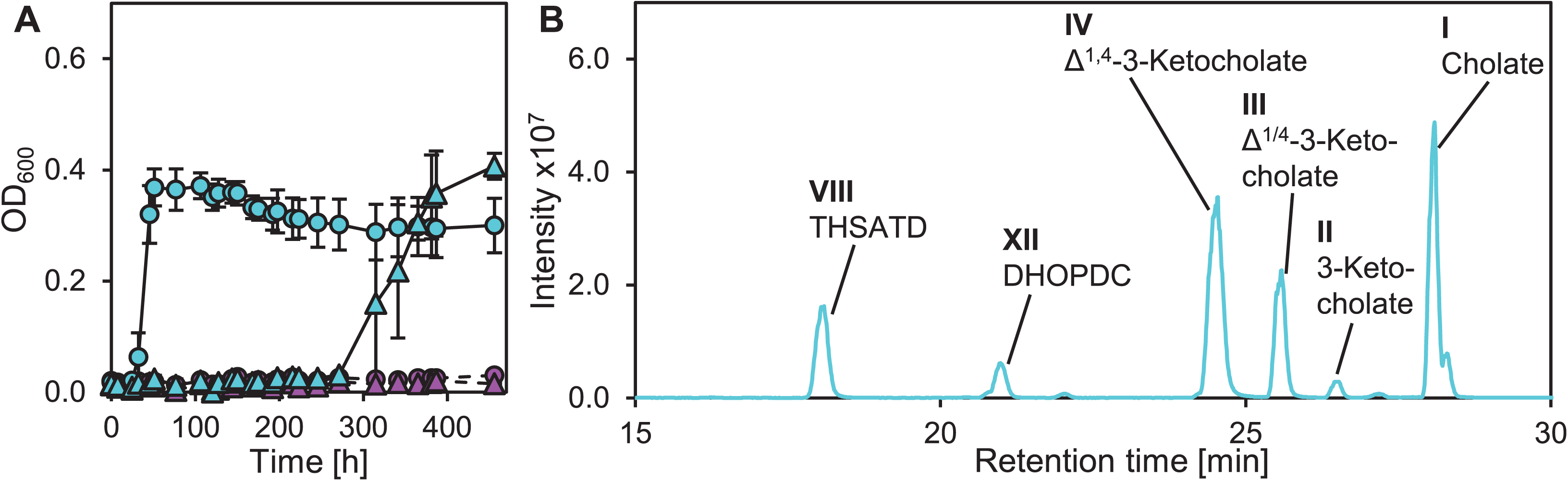
Heterologous complementation of *P. stutzeri* Chol1 Δ*scd1A* and Δ*scd1B* with *scd4AB.* (A) Growth of cross-complementation strains *P. stutzeri* Chol1 Δ*scd1A* pBBR1MCS-5::*scd4AB* (turquois circles) and Δ*scd1B* pBBR1MCS-5::*scd4AB* (turquois triangles) as well as empty vector controls *P. stutzeri* Chol1 Δ*scd1A* pBBR1MCS-5 (red circles) and Δ*scd1B* pBBR1MCS-5 (red triangles) with 1 mM cholate. Error bars indicate standard deviations and may not be visible if too small (n=3). (B) Transient accumulation of various intermediates in the supernatant of a *P. stutzeri* Chol1 Δ*scd1A* pBBR1MCS-5::*scd4AB* culture with cholate at an OD_600_ of about 0.13. MS base peak chromatogram in negative ion mode. Intermediates were identified according to retention time, UV absorbance, and mass.

We also tested whether Nov2c221 and Nov2c222 might be able to take over the function of the second ACAD pair in *P. stutzeri* Chol1 by heterologous expression in *P. stutzeri* Chol1 R1, which is a transposon mutant defective in Scd2AB (16). However, expression of *nov2c221* and *nov2c222* in *P. stutzeri* Chol1 R1 could not restore growth and degradation of cholate (data not shown).

Together these results strongly indicate, that Nov2c221 and Nov2c222 form an ACAD that introduces a double bond in the C_5_-side chain of cholate at C22 and were therefore renamed Scd4A and Scd4B, respectively, for **s**teroid-acyl-**C**oA-**d**ehydrogenase.

### Strain Chol11 Δ*scd4A* transforms bile salts to hydroxylated metabolites with complete side chain that can be degraded by the wild type

To explore the metabolic capacities of *Sphingobium* sp. strain Chol11 Δ*scd4A*, the strain was incubated for prolonged time periods with different bile salts and glucose as an additional carbon source. Apart from the trihydroxy bile salt cholate (I in Fig 1), the dihydroxy bile salts deoxycholate (XVI in Fig 4) and chenodeoxycholate (XV) as well as the monohydroxy bile salt lithocholate (XIV) were used, which can all be completely degraded by the strain Chol11 wt (28). After incubation for two weeks, strain Chol11 Δ*scd4A* had transformed all bile salts to several metabolites that were not degraded upon further incubation (Fig 4). Four (P1-P4) out of five metabolites from the transformation of lithocholate and chenodeoxycholate overlapped (Fig 4B), as well as those from transformation of cholate and deoxycholate (P6-P8, Fig 4C). In general, the masses of these metabolites indicated that the side chains of the bile salts remained complete; none of these metabolites had an absorption shoulder around 210 nm that is indicative of the additional double bond at C22 (12, 22). Interestingly, mass spectra indicated that some of the metabolites were hydroxylated. To investigate whether these metabolites were true intermediates of bile-salt degradation or dead-end products, filter-sterilized culture supernatants of lithocholate- and cholate-grown cells of strain Chol11 Δ*scd4A* were mixed with fresh medium and used for growth experiments with strain Chol11 wt. While P1, P3, P5, P6, and P8 were completely degraded by strain Chol11 wt, P2 and P7 remained (Fig S4A+B).

**Fig 4.**
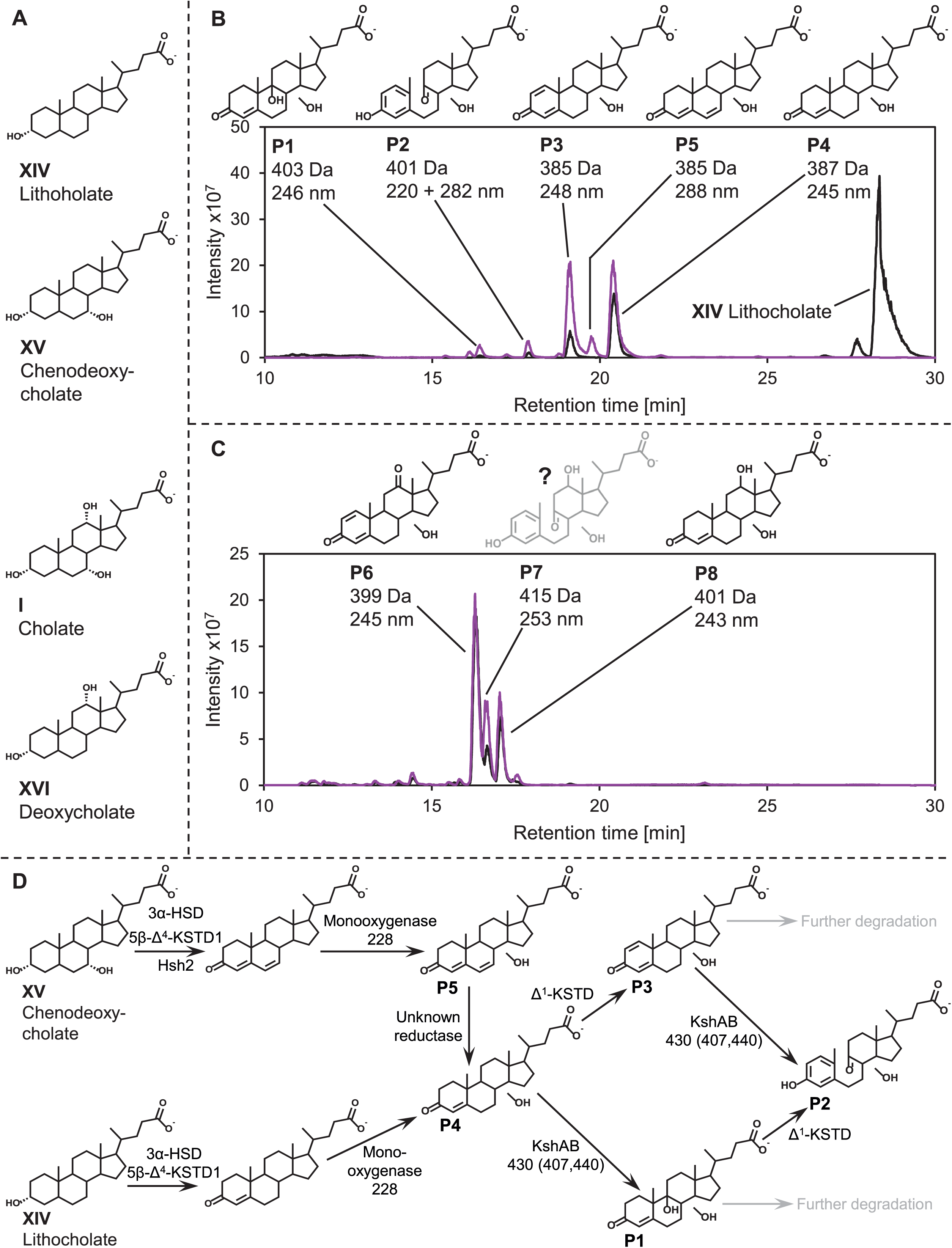
Transformation of bile salts by *Sphingobium* sp. strain Chol11 Δ*scd4A.* (A) Structures of the tested bile salts. (B) Accumulation of several dead-end products (P1-P5) by biotransformation of 12-deoxy bile salts chenodeoxycholate (red) and lithocholate (black) by strain Chol11 Δ*scd4A.* (C) Accumulation of several dead-end products (P6-P8) by biotransformation of 12-hydroxy bile salts cholate (black) and deoxycholate (red) by strain Chol11 Δ*scd4A*. MS base peak chromatogram in negative ion mode. Masses are indicated for the respective deprotonated acids. Structure suggestions are based on retention time, UV absorbance, and mass. (D) Potential pathway for transformation of chenodeoxycholate (XV) and lithocholate (XIV) to products P1-5 by strain Chol11 Δ*scd4A.* 3α-HSD: 3α-hydroxysteroid dehydrogenase (e.g. Nov2c6), 5β-Δ^4^-KSTD1: 5β-Δ^4^-ketosteroid dehydrogenase Nov2c19, Hsh2: Nov2c400, Δ^1^-KSTD: Δ^1^-ketosteroid dehydrogenase. Locus tags of monooxygenases are given in parentheses (e.g. 228 for Nov2c228). Grey: P1 and P3 can be completely degraded by strain Chol11.

### Two of the degradable steroid metabolites can be activated by CoA-ligase SclA

To obtain further evidence that the degradable metabolites are true intermediates of the metabolic pathway, it was tested, if they could be activated by steroid CoA-ligase SclA from strain Chol11 (12). In assays with cell free extract of *E. coli* MG1655 expressing *sclA*, two compounds with masses of 1134.5 Da and 1136.5 Da and characteristic absorption maxima at about 250 nm were formed when CoA and ATP were present (Fig 5A). The mass of 1136.5 Da indicates, that this compound is the CoA-ester of P4 (387 Da of P4 + 767.5 Da of CoA – 18 Da H_2_O which is removed for thioester formation) (Fig 5B) and the mass of 1134.5 Da accordingly indicates, that this compound is the CoA-ester of P3 (385 Da of P3 + 757.5 Da of CoA – 18 Da H_2_O) (Fig 5C). The UV-absorption spectra of these compounds are also indicative of CoA-esters (16). Both CoA-esters were not formed when the cell extract was omitted and when cell extracts of *E. coli* empty vector controls were used. This indicates that the activation is catalyzed by SclA. Minor formation of these CoA-esters in assays without either CoA or ATP could be due to residual CoA and ATP in the cell-free extracts.

**Fig 5.**
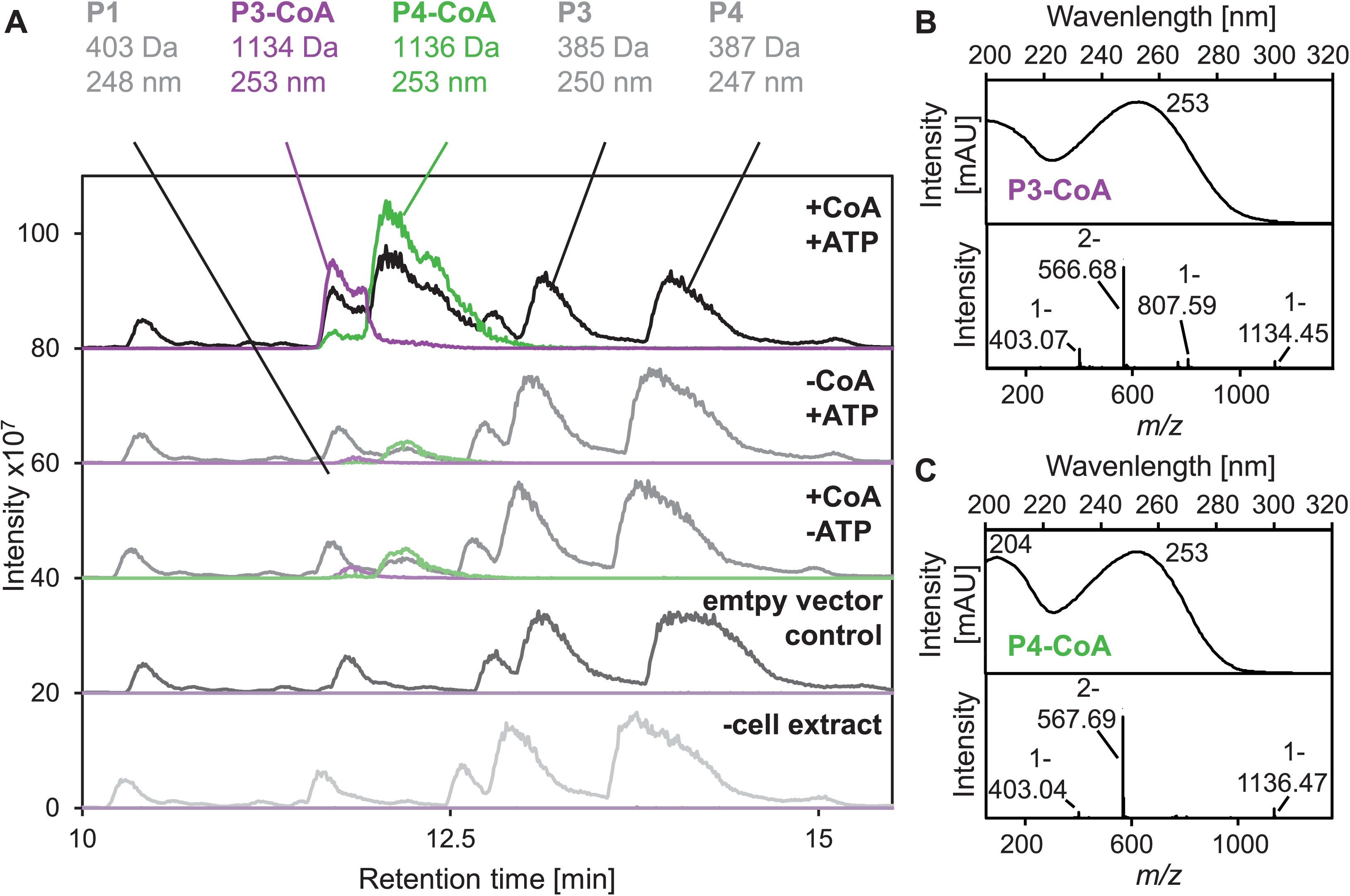
(A) CoA-activation of the metabolites P1-P4 produced from lithocholate by *Sphingobium* sp. strain Chol11 Δ*scd4AB* by cell-free extract of *E. coli* MG1655 pBBR1MCS-5*::sclA* after 4 h at 30 °C. MgCl_2_ was added in all assays. MS chromatograms in negative ion mode of the CoA-activation assay and controls as indicated: Black and grey, base peak chromatogram; red, extracted ion chromatogram for *m/z* = 566.7 in negative mode (dominant *m/z* value of P3-CoA: [M+H]^-2^ = 566.7 Da); green, extracted ion chromatogram for *m/z* = 567.7 in negative mode (dominant *m/z* value of P3-CoA: [M+H]^-2^ = 567.7 Da). Steroid compounds were assigned to structures by retention time, UV absorbance, and mass. Chromatograms are shown with offset in intensity for better overview. Masses calculated from negative mode MS measurements and absorption maxima are given. Masses are indicated for the respective deprotonated acids. P2 was not detected in these measurements, probably due to insufficient separation with the altered method. (B+C) UV- and MS-spectrum of the CoA-activated P3 (B) and P4 (C) produced in the enzyme assays, retention times 11.7 min and 12.1 min, respectively.

### The metabolites produced by strain Chol11 Δ*scd4A* are mono- or dihydroxylated and have Δ^1,4^-structures or cleaved B-rings

As it was not possible to produce the unknown metabolites and dead-end products in sufficient amounts in the required purity for NMR analyses, information on the structure could only be inferred indirectly. Strain Chol11 Δ*scd4A* transformed both lithocholate and chenodeoxycholate to the apparently identical products P1, P3, and P4 (Fig 4B). According to their absorption maxima at approximately 245 nm, products P1, P3, and P4 had Δ^4^- or Δ^1,4^-3-keto structures of the A-ring (Fig S5); such compounds have previously been detected in supernatants of strain Chol11 (11, 27, 28). According to their masses, P3 (385 Da) and P4 (387 Da) have an additional hydroxy group at an unknown position compared to Δ^1,4^- and Δ^4^-3-ketolithocholate, respectively. For finding the position and stereochemistry of this hydroxy group, P4 was compared to Δ^4^-3-ketochenodeoxycholate, Δ^4^-3-ketohyodeoxycholate, and Δ^4^-3-ketoursodeoxycholate with hydroxy groups in 7α-, 6α- or 7β-position, respectively (Fig S6A). These reference compounds were produced from the parent bile salts with mutants from our strain collection as described in the methods section. Although all substances had similar retention times ranging from 19 to 20 min, the retention time of P4 was nearly identical to Δ^4^-3-ketochenodeoxycholate. As Δ^4^-3-ketochenodeoxycholate is a substrate for Hsh2 (28), we tested, if P4 could be transformed by Hsh2. However, no formation of the respective Δ^4,6^-3-keto product could be observed in enzyme assays with P4 and Hsh2 (Fig S6B). Thus, the hydroxy group of P4 was apparently in a different position that could not be defined yet. The masses of the other two metabolites P1 (403 Da) and P2 (401 Da) indicate two hydroxy groups compared to Δ^4^- and Δ^1,4^-3-ketolithocholate, respectively. While the absorption spectrum of P1 indicates a Δ^4^-3-keto structure, the absorption spectrum of P2, which is very similar to THSATD (VIII in Fig 1), indicates a 9,10-*seco* structure that could plausibly be caused by hydroxylation at C9 (23); this *seco*-steroid with C5-side chain could apparently not be degraded by strain Chol11 wt.

P5 was not found in cultures of strain Chol11 Δ*scd4A* grown in the presence of lithocholate but was unique to growth in presence of chenodeoxycholate. The UV-spectrum of P5 was indicative of a Δ^4,6^-3-keto structure of the steroid skeleton, which agrees with the mutant strain still exhibiting 7α-dehydratase activity. As the mass of P5 indicated an additional hydroxy group, it can most likely be excluded that this hydroxy group is at C7 but rather at the same position as in P3 and P4.

Cholate and deoxycholate were also transformed to identical products (Fig 4C. Assuming that the 12-hydroxy bile salts cholate and deoxycholate are transformed like chenodeoxycholate and lithocholate, P6 (399 Da, 245 nm) and P8 (401 Da, 243 nm) most probably are the 12-hydroxy derivatives of P3 and P5, respectively, according to their masses and absorption spectra (Fig S5). The absorption maximum of P7 at 253 nm is not typical for steroid compounds and, consequently, no structure could be assigned to this compound. However, its mass indicated that it was hydroxylated twice compared to the precursor cholate.

### Strain Chol11 lacks candidate genes for further degradation of the side chain as known from strain Chol1

The next step in steroid side-chain degradation during both aldolytic and thiolytic degradation in *P. stutzeri* Chol1 and *R. jostii* RHA1 is the addition of water to the double bond to gain a β-hydroxy group (17, 30). The only candidates for this reaction that could be found in genomic and proteomic analyses (12, 13 (preprint)), were *nov2c219* and *nov2c220* in the side-chain degradation gene cluster (Fig S1). However, the deletion mutant strain Chol11 Δ*nov2c219-220* did not differ significantly from the wild type during growth with cholate (Fig S7A). Cross-complementation with plasmid-encoded *nov2c219* and *nov2c220* could not restore growth of the deletion mutant *P. stutzeri* Chol1 Δ*shy*, which lacks the hydratase necessary for C_5_ side-chain degradation. Furthermore, deletion of the adjacent gene *nov2c218*, which encodes a short chain reductase with 31 % similarity to the 7α-hydroxysteroid dehydrogenase of *E. coli* (31) did also not cause a phenotype during growth with cholate (Fig S7B).

### Five potential steroid hydroxylating Rieske monooxygenases and homologs to KshA are encoded in the genome of strain Chol11

*Sphingobium* sp. strain Chol11 Δ*scd4A* transformed bile salts to *seco*-steroids with cleaved B-ring as well as further hydroxylated bile-salt derivatives indicating the activity of monooxygenases for 9α-hydroxylation as well as for hydroxylation at other positions. Therefore, we aimed at elucidating the function of putative steroid monooxygenases of strain Chol11 as a next step. One P450 monooxygenase is encoded near one steroid-degradation gene cluster of strain Chol11 but was not detected in any cells in proteomic analyses (13 (preprint)). Additionally, strain Chol11 encodes five homologs of the oxygenase component KshA of the ring cleaving 9α-monooxygenase KshAB, which belong to the Rieske monooxygenases (Tab 2) (13 (preprint)). Interestingly, Nov2c66, Nov2c407, Nov2c430 and Nov2c440 had high similarities of >40 % among themselves, while Nov2c228 is more different with <30 % identity to the others (Table 2, Fig S8). Nov2c407, Nov2c430 and Nov2c440 were produced specifically during degradation of steroids whereas Nov2c228 was only produced during degradation of side-chain bearing steroids (13 (preprint)). For the reductase component KshB, only one putative homolog could be found (Table 2), which had less than 20% identity to KshB from *R. jostii* RHA1.

**Table 2.** KshA and KshB homologs from *Sphingobium* sp. strain Chol11 and *P. stutzeri* Chol1 compared to KshA1 and KshB from *R. rhodochrous* DSM43269. KshA1 is the KshA homolog from *R. rhodochrous* DSM43269 involved in degradation of bile salts.

| **KshA** |  |  |  |  |
| --- | --- | --- | --- | --- |
| Name | Identity [%] to |  |  | RefSeq ID |
|  | KshA1<sub>DSM43269</sub> | KshA<sub>Chol1</sub> | Nov2c407 |  |
| Nov2c66 | 26 | 32 | 58 | WP_097093404.1 |
| Nov2c228 | 23 | 27 | 25 | WP_097093548.1 |
| Nov2c407 | 29 | 31 | 100 | WP_097092973.1 |
| Nov2c430 | 28 | 31 | 45 | WP_097092990.1 |
| Nov2c440 | 29 | 31 | 99 | WP_097093001.1 |
| C211_11582/KshA<sub>Chol1</sub> | 38 | 100 |  | WP_008568691.1 |
| **KshB** |  |  |  |  |
| Name | Identity [%] to |  | RefSeq ID |  |
|  | KshB<sub>DSM43269</sub> | KshB<sub>Chol1</sub> |  |  |
| Novbp123 | 18 | 17 | WP_013039106.1 |  |
| C211_11317/KshB<sub>Chol1</sub> | 50 | 100 | WP_008568639.1 |  |

As the multiplicity of KshA homologs would most probably necessitate multiple deletions for completely abolishing KshA activity, we conducted a heterologous expression. So far, no KshAB from Proteobacteria has been characterized, so we first aimed at constructing a *kshA* deletion mutant in *P. stutzeri* Chol1 as a heterologous expression platform.

### *P. stutzeri* Chol1 has a single KshAB that can functionally be expressed in ***Escherichia coli***

In *P. stutzeri* Chol1, only one homologous protein for each KshA and KshB can be found. These are C211_11582 (RefSeq-ID WP_008568691.1) and C211_11317 (WP_008568639.1) with 38% and 50% identity to KshA1 and KshB from *Rhodococcus rhodochrous* DSM43269, respectively (Table 2, Fig S8). Both respective genes are located in a steroid degradation cluster (17, 20, 32). A deletion mutant of *kshA_Chol1_* was constructed that showed strongly decreased growth with cholate (I in Fig 1) (Fig 6A) and accumulated a single metabolite that could be identified as 12β-DHADD (VI) (Fig 6B) according to its retention time, mass (316 Da), and UV-spectrum (maximum at 245 nm). With chenodeoxycholate (XV in Fig 4), deoxycholate (XVI), and lithocholate (XIV) *P. stutzeri* Chol1 Δ*kshA* had the same phenotype and accumulated the respective ADDs derived from these bile salts (Fig S9). Growth could be restored by plasmid-borne *kshA_Chol1_* (Fig 7A). In the next step, *kshA* and *kshB* were co-expressed in *E. coli* MG1655. In cell suspensions supplied with androsta-1,4-diene-3,17-dione (ADD, XIX in Fig S9), *E. coli* MG1655 pBBR1MCS-5::*kshA_Chol1_-kshB_Chol1_* was able to cleave the substrate and form the 9,10-*seco*-steroid 3-hydroxy-9,10-*seco-*androsta-1,3,5-triene-9,17-dione (HSATD, XX in Fig 8) (Fig 8A). These reactions were not observed in suspensions of the vector control (Fig 8B), indicating that this is a functional platform for observing the activities of KshAB complexes (16, 17, 22). To further explore the substrate spectrum of KshAB_Chol1_, a variety of metabolites with Δ^1,4^-3-keto structure, which had either no or different side chains, were tested in this platform. KshAB_Chol1_ was also able to hydroxylate and cleave all tested Δ^1,4^-metabolites without side chain irrespective of the hydroxylation pattern of the steroid skeleton (Fig S10).

**Fig 6.**
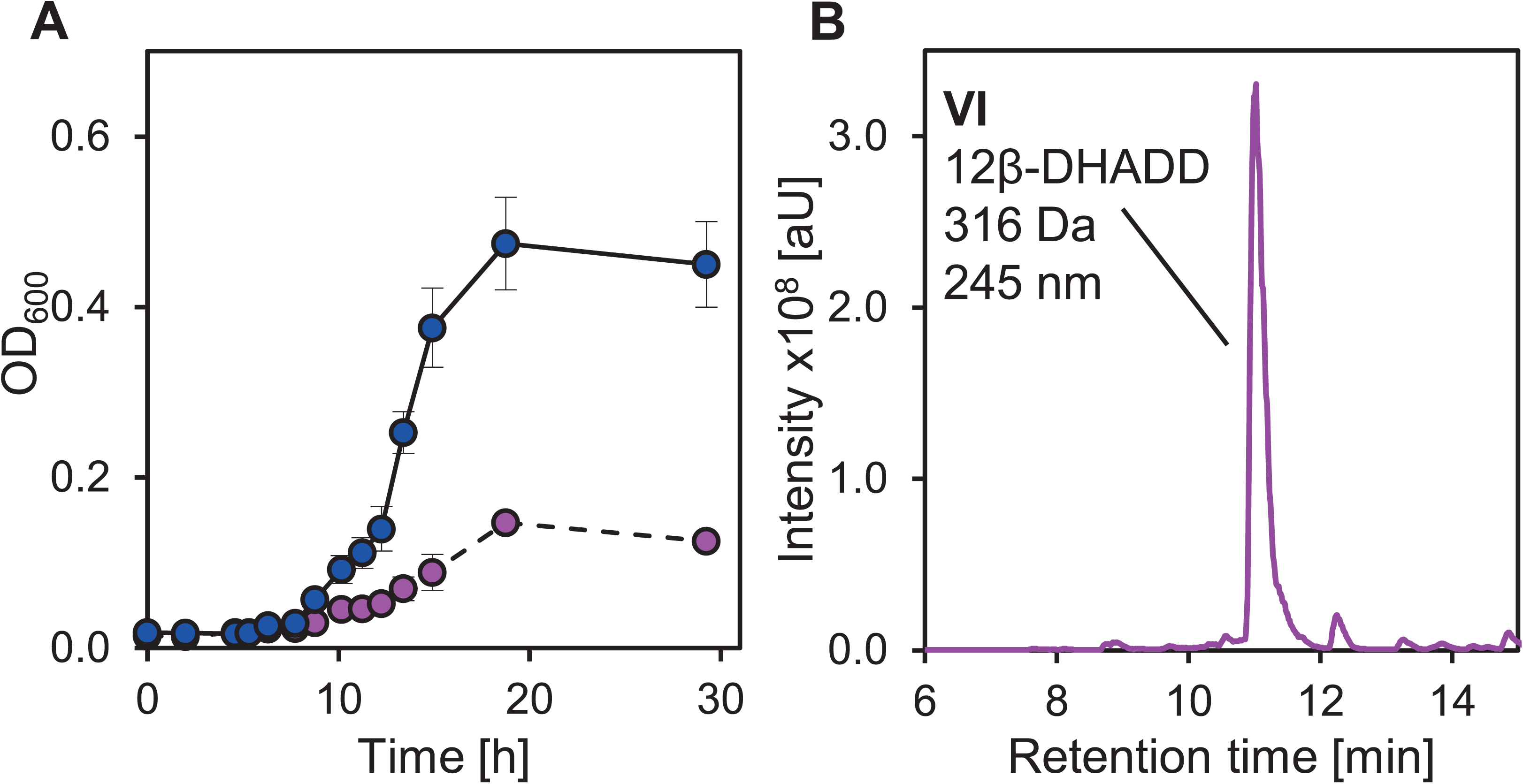
Phenotype of *P. stutzeri* Chol1 Δ*kshA.* (A) Growth of *P. stutzeri* Chol1 Δ*kshA* (dashed line, red) and wt (solid line, blue) with 1 mM cholate. Error bars indicate standard deviations and may not be visible if too small (n=3). (B) Accumulation of 12β-DHADD (VI in Fig 1) as dead-end metabolite in the supernatant of *P. stutzeri* Chol1 Δ*kshA* grown with cholate for about 30 h. MS base peak chromatogram in positive ion mode.

**Fig 7.**
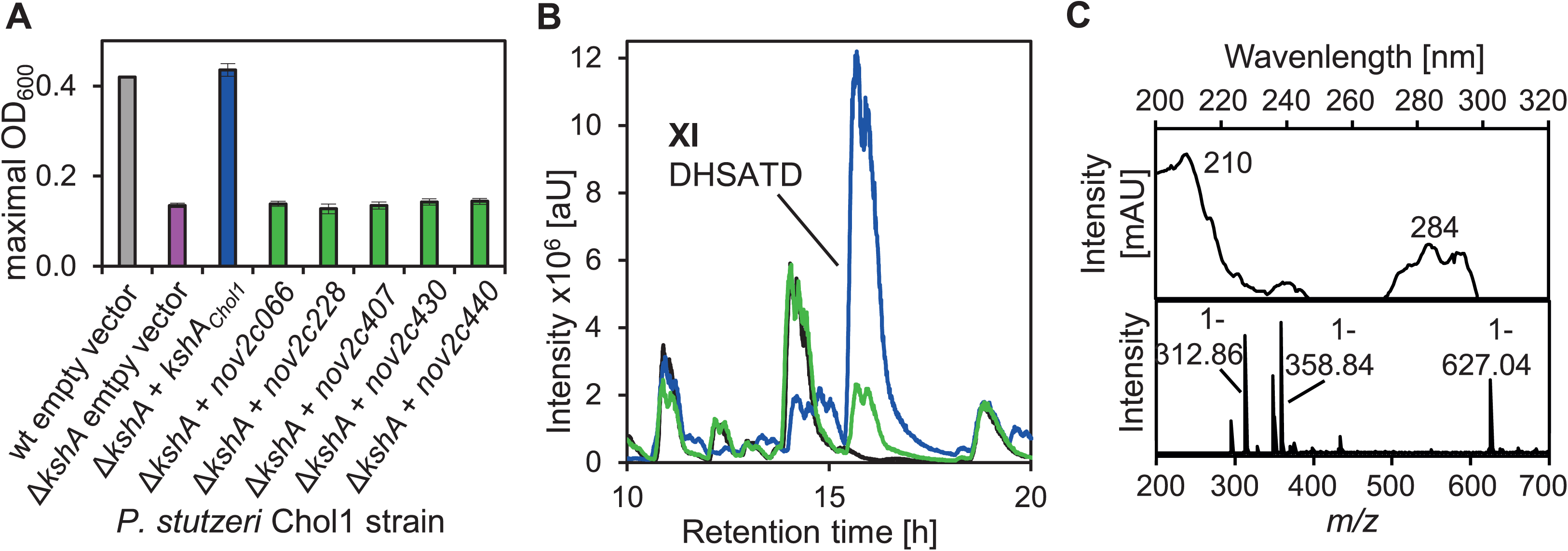
Heterologous complementation of *P. stutzeri* Chol1 Δ*kshA.* (A) Maximal OD_600_ reached by *P. stutzeri* Chol1 wt and Δ*kshA* complemented with the given genes on vector pBBR1MCS-5 when grown with 1 mM cholate. (B) Transformation of HOCDA (V in Fig 1) to DHSATD (XI) by *P. stutzeri* Chol1 Δ*kshA* expressing different genes on plasmid pBBR1MCS-5. HPLC-MS base peak chromatograms in negative mode of the supernatants of cell suspensions of *P. stutzeri* Chol1 pBBR1MCS-5::*nov2c430* (green), pBBR1MCS-5::*kshA* (blue), and Δ*kshA* pBBR1MCS-5 (empty vector control, black) supplemented with HOCDA and incubated for 6 days. (C) UV- and MS-spectrum of DHSATD produced by *P. stutzeri* Chol1 Δ*kshA* pBBR1MCS-5::*nov2c430*.

**Fig 8.**
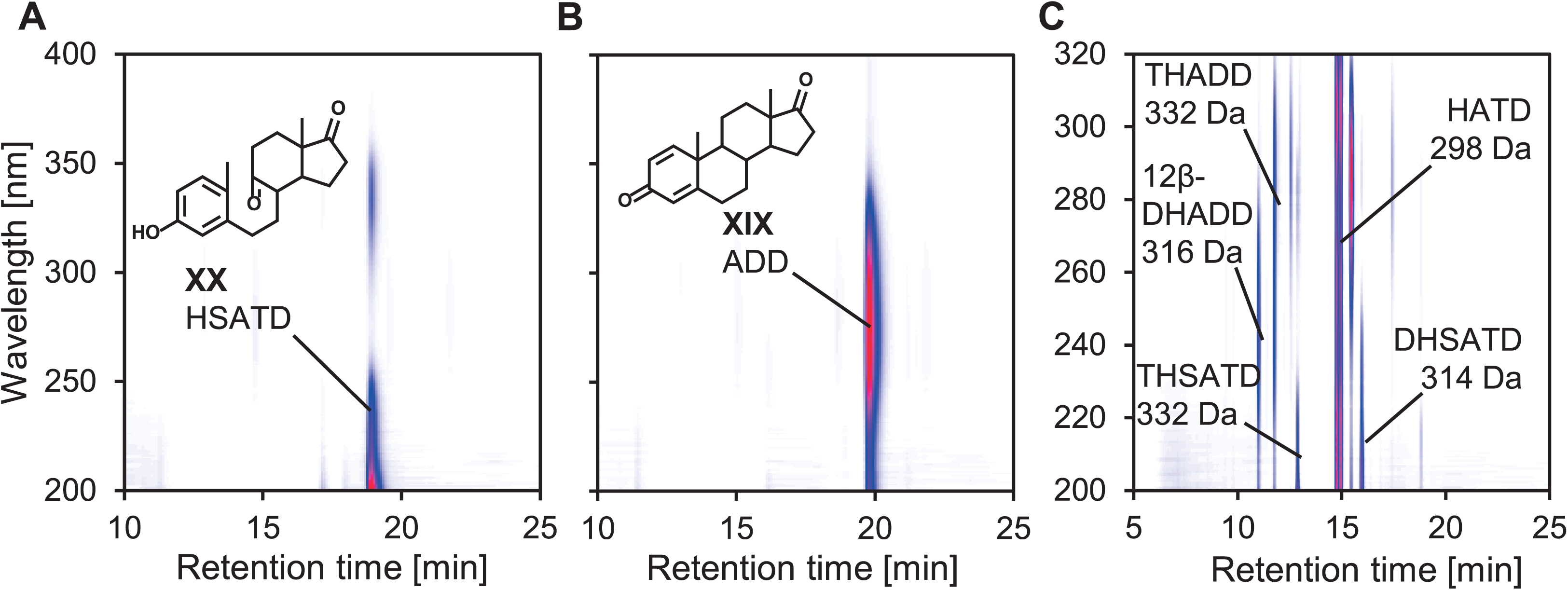
Transformation of steroid compounds by *E. coli* MG1655 expressing *kshAB_Chol1_.* (A) Transformation of ADD (XIX) to the respective 9,10-*seco*-steroid 3-hydroxy-9,10-*seco*-androsta-1,3,5(10)-triene-9,17-dione (HSATD, XX) by *E. coli* MG1655 pBBR1MCS-5::*kshAB_Chol1_.* (B) No transformation of ADD by empty vector control *E. coli* MG1655 pBBR1MCS-5. (C) Transformation of HATD (X in Fig 1) and 12β-DHADD (VI) by *E. coli* MG1655 pBBR1MCS-5::*kshAB_Chol1_* to the *seco-*steroids THSATD (VIII) and DHSATD (XI) and the side-product 1,2,12-trihydroxy-androsta- 4,6-diene-3,17-dione (THADD, XXIII in Fig S13). 3D-UV chromatograms of supernatants of cell suspensions of the respective *E. coli* strain incubated for 4 days with ADD. Red indicates highest absorption. Steroid compounds were identified by retention time, UV absorbance, and mass.

### KshAB_Chol1_ is active with a metabolite from the Δ^4,6^-variant and produces a dihydroxylated dead-end metabolite

*P. stutzeri* Chol1 has already been shown to transform HOCDA (V in Fig 1) to HATD (X) that is further converted to the 9,10-*seco* steroid DHSATD (XI) indicating functional 9α-hydroxylation with substrates from the Δ^4,6^-variant (11). To investigate whether this reaction is also catalyzed by KshAB in *P.stutzeri* Chol1, we supplied the *kshA* deletion mutant with HOCDA. *P. stutzeri* Chol1 Δ*kshA* with the empty vector pBBR1MCS-5 transformed HOCDA only to HATD (X in Fig 1) (Fig S11B), whereas the complemented mutant expressing *kshA*_Chol1_ like the wild type *P. stutzeri* Chol1 (Fig S11A+C) formed DHSATD, which cannot be further degraded by *P. stutzeri* Chol1 (11). Some THSATD (VIII) produced by *P. stutzeri* Chol1 wt and the mutant complemented with *kshAB_Chol1_* as well as 12β-DHADD (VI) produced by the empty vector control were likely derived from a Δ^4^-3-ketocholate contamination in the HOCDA stock solution.

In the next step, we supplied HATD to the *E. coli* expression platform. *E. coli* pBBR1MCS-5::*kshAB_Chol1_* transformed HATD (X) into DHSATD (XI) (Fig 8C in contrast to a control in Fig S12). Some THSATD (VIII) found in these supernatants is most likely derived from a contamination of the HATD stock solution with 12β-DHADD (VI).

During transformation of HOCDA by *P. stutzeri* Chol1, we previously observed the formation of a dihydroxylated metabolite 1,2,12-trihydroxy-androsta-4,6-diene-3,17-dione (THADD, XXIII in Fig S13, Fig S11A+C) (11), which was formed as a further dead-end metabolite in addition to DHSATD. In our transformation experiment with the *kshA_Chol1_* deletion mutant and HOCDA as substrate, no THADD was formed (Fig S11B). When HATD was supplied to *E. coli* pBBR1MCS-5::*kshAB_Chol1_* the formation of some THADD could be shown according to its retention time, mass (332 Da), and UV-spectrum (maximum at 290 nm) (Fig 8C, Fig S13). These experiments showed that KshAB_Chol1_ is not only able to catalyse 9α-hydroxylation of a substrate from the Δ^4,6^-variant but also to catalyze a side reaction on the A-ring.

### Three out of five KshA homologs from *Sphingobium* sp. strain Chol11 have B-ring cleaving activity in *P. stutzeri* Chol1 with HOCDA as substrate

After KshA_Chol1_ had been characterized, the respective homologues from strain Chol11 were studied. First, all five *kshA* homologs from *Sphingobium* sp. strain Chol11 were expressed in *P. stutzeri* Chol1 Δ*kshA* to check for cross-complementation. However, none of them was able to replace natural *kshA*_Chol1_ during growth with cholate (Fig 7A) and all cross-complemented strains accumulated 12β-DHADD as single product from cholate.

As we proposed that Δ^4,6^-intermediates are substrates for B-ring cleavage in strain Chol11, we next supplied the Δ^4,6^ intermediate HOCDA (V in Fig 1) as an alternative substrate (Fig S11, Table 3). Strains expressing *nov2c407, nov2c430,* or *nov2c440* produced DHSATD (XI in Fig 1) (shown for *nov2c430* in Fig 7B+C and Fig S11F, shown for *nov2c407* and *nov2c440* in Fig S11G-J). This conversion was much less efficient than with the innate KshA-homolog of *P. stutzeri* Chol1 (see above; Fig S11C). Nevertheless, DHSATD formation clearly indicated steroid 9α-monooxygenase activity of Nov2c407, Nov2c430, and Nov2c440 in *P. stutzeri* Chol1 leading to B-ring cleavage.

**Table 3.**
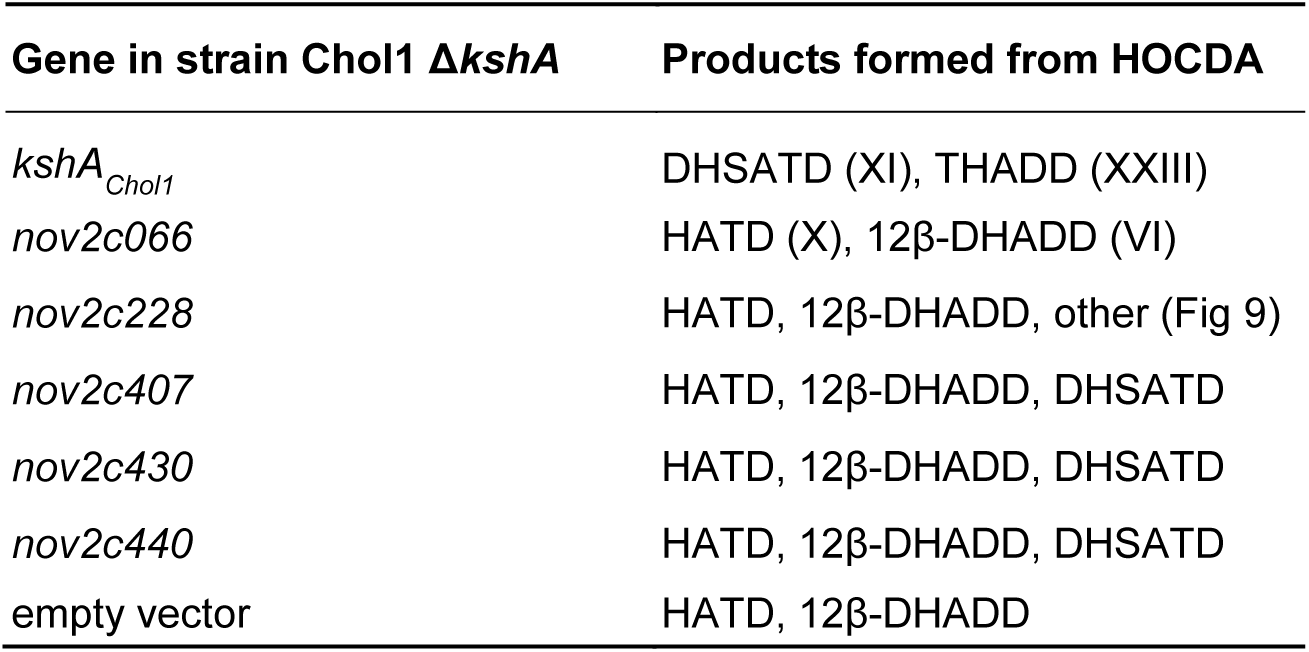
Metabolites produced by transformation of HOCDA (V) by *P. stutzeri* Chol1 Δ*kshA* complemented with the given genes on vector pBBR1MCS-5 after 6 days as determined by HPLC-MS and identified by retention time, mass, and UV-spectrum. THADD, 1,2,12-trihydroxy-androsta-4,6-diene-3,12-dione (XXIII in Fig S13).

| Gene in strain Chol1 $\Delta kshA$ | Products formed from HOCDA |
| --- | --- |
| *kshA*<sub>Chol1</sub> | DHSATD (XI), THADD (XXIII) |
| *nov2c066* | HATD (X), 12β-DHADD (VI) |
| *nov2c228* | HATD, 12β-DHADD, other (Fig 9) |
| *nov2c407* | HATD, 12β-DHADD, DHSATD |
| *nov2c430* | HATD, 12β-DHADD, DHSATD |
| *nov2c440* | HATD, 12β-DHADD, DHSATD |
| empty vector | HATD, 12β-DHADD |

These DHSATD-forming *kshA* homologs from strain Chol11 were also expressed in *E. coli* either in combination with KshB_Chol1_ or with the putative KshB homolog Novbp123 from strain Chol11. For providing a diverse set of substrates, cell suspensions were incubated with several different steroid compounds (Fig S10). Neither hydroxylation nor ring cleavage of any of the substrates including the presumptive physiological substrate HATD could be observed for any putative KshA_Chol11_ with either KshB homolog by HPLC-MS measurements (data not shown). Notably, KshA_Chol1_ was only active with KshB from strain Chol1 and not with Novbp123 (Fig S10).

### Rieske monooxygenase Nov2c228 has steroid-hydroxylating activity in *P. stutzeri* Chol1

*P. stutzeri* Chol1 Δ*kshA* with Nov2c228 did not catalyze B-ring cleavage but transformed HOCDA to four products (P9-11, XXI), that were not produced by the empty vector control or the wild type (Fig 9, Fig S14). Interestingly, two of the products (P9 and P10) could be assigned to structures with an additional hydroxylation at an unknown position, potentially similar to the mono-hydroxylated dead-end products of strain Chol11 Δ*scd4A*. P10 has a Δ^1,4,6^-3-keto structure as found in HATD (X in Fig 1) according to its absorption spectrum, and its mass indicated a C_3_-side chain and an additional hydroxy group at a so-far unknown position. In contrast, P9 has a Δ^1,4^-3-keto structure, a C_3_-side chain and an additional hydroxy group according to its absorption spectrum and mass, and therefore may be derived from a Δ^4^-3-ketocholate contamination of the HOCDA stock solution. Two more products did not have additional hydroxylations but most probably were 12-hydroxy-3-oxo-pregna-1,4,6-triene-carboxylate (HOPTC, XXI in Fig 9) with a C_3_ carboxylic side chain, and Δ^1,4,6^-3,12-diketocholate according to their masses and absorption spectra. In the *E. coli* system, Nov2c228 did not show activity towards any of the tested substrates with either KshB homolog.

**Fig 9.**
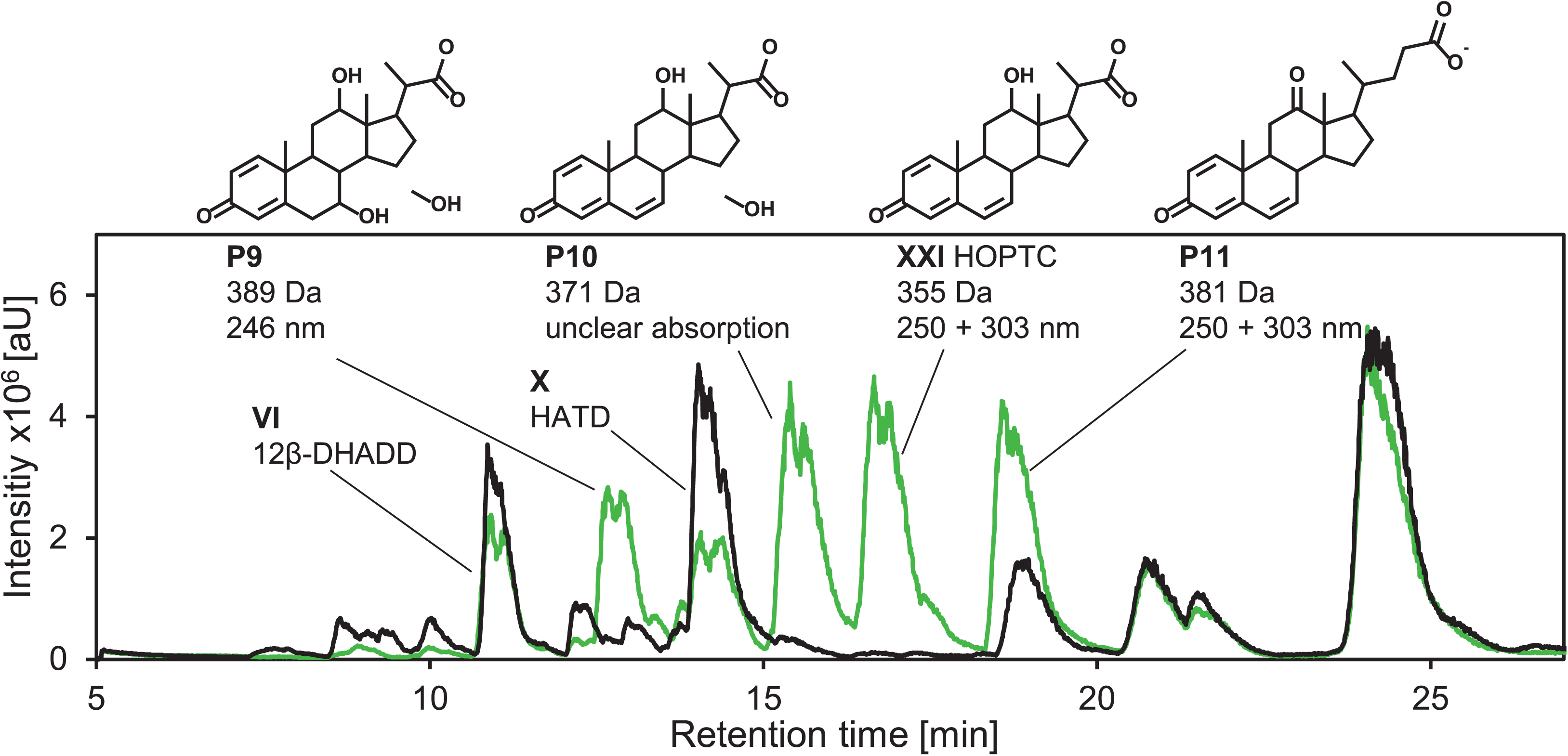
Transformation of HOCDA (V in Fig 1) by *P. stutzeri* Chol1 Δ*kshA* pBBR1MCS-5::*nov2c228* (green) and empty vector control *P. stutzeri* Chol1 Δ*kshA* pBBR1MCS-5 (black). Base peak MS chromatograms in negative ion mode of supernatants of cell suspensions of the respective strain incubated for 6 days with HOCDA. Identification of steroid compounds and structures suggestions are based on retention time, UV absorbance, and mass. Masses calculated from negative mode MS measurements and absorption maxima are given. Masses are indicated for the respective deprotonated acids. XXI, 12-hydroxy-3-oxo-pregna-1,4,6-triene-carboxylate (HOPTC). The substances eluting at 21 min, 21.5 min, and 24.5 min cannot be assigned to any structures and are most probably no steroidal compounds according to their UV- and MS-spectra.

*P. stutzeri* Chol1 Δ*kshA* expressing the fifth KshA homolog Nov2c66 did not produce any hydroxylated metabolites (Fig S11D), and no activity of Nov2c66 could be observed in the *E. coli* system.

## Discussion

After genomic (12) and proteomic studies (13 (preprint)) have already suggested that the degradation of the bile-salt side chain in *Sphingobium* sp. strain Chol11 proceeds differently from the reaction found in Pseudomonads, Comamonads, and Actinobacteria, our study provides further functional evidence in comparative studies with strain Chol11 in contrast to *P. stutzeri* Chol1. Our results reveal that the ACAD reaction for introducing a Δ^22^-double bond catalyzed by Scd4AB is necessary for side-chain degradation, which is still equivalent to the mechanism observed in the model organisms mentioned above. Strain Chol11 Δ*scd4A* is the first deletion mutant of strain Chol11 that is not able to grow with cholate anymore (12, 28). However, the next steps of side-chain degradation are apparently different and most probably involve a hydroxylation step that might plausibly be catalyzed by the Rieske monooxygenase Noc2c228. As a further result regarding bile-salt degradation in strain Chol11, we obtained evidence that B-ring cleavage is also proceeding via 9α-hydroxylation leading to 9,10-seco structures with DHSATD (XI in Fig 1) as intermediate because the Rieske monooxygenases Nov2c407, Nov2c430, and Nov2c440 catalyzed the formation of DHSATD from HOCDA (V) in the *kshA*-deletion mutant of strain Chol1 as heterologous host.

The physiological role of Scd4AB as an ACAD for the introduction of a double bond at C22 was concluded from the functional chemical complementation of the growth-defective of the respective deletion mutant with DHOCTO (XIII in Fig 2) that already had this double bond. Furthermore, heterologous expression of Scd1AB and Scd4AB in the individual ACAD deletion mutants of strain Chol11 and *P. stutzeri* Chol1, respectively, also enabled genetic complementation. Obviously, the complexes Scd4AB from strain Chol11 and Scd1AB from *P. stutzeri* Chol1 were interchangeable. However, both innate subunits were needed to form an active ACAD complex indicating that the ACADs are too different to form promiscuous complexes. Furthermore, the strain Chol11 Δ*scd4A* deletion mutant complemented with *scd1AB* grew with cholate without any difference to the wild type while the strain Chol1 Δ*scd1A* and Δ*scd1B* deletion mutants, respectively, complemented with *scd4AB* showed strongly extended lag phases of varying length as well as reduced growth. These differing phenotypes could plausibly be explained by the fact that the CoA-ester of HOCDA (V in Fig 1) with a Δ^4,6^-structure of the steroid skeleton is the presumptive native substrate of Scd4AB while Scd1AB oxidizes the CoA-ester of Δ^1,4^-3-ketocholate (20). From previous studies it is known that HOCDA can be readily converted by the side-chain degrading enzymes of strain Chol1 (11, 28). Scd4AB might be less active with steroid-acyl-CoA substrates that have a Δ^1,4^-structure of the steroid skeleton, and might require a Δ^4,6^-substrate, which was not supplied in strain Chol1, for its full activity. Thus, the differences between Scd1AB and Scd4AB indicated by the phylogenetic analyses, that may hint at a different phylogenetic origin of this side-chain degradation mechanism, were also reflected by this functional analysis.

The physiological role of the involvement of a hydroxylation for side-chain degradation is based on the interconnection of multiple independent results. First, strain Chol11 Δ*scd4A* transformed cholate, deoxycholate, chenodeoxycholate, and lithocholate to several mono- and dihydroxylated steroid compounds with complete side chain. It is known from *P. stutzeri* Chol1 that hydroxylations may occur as side reactions when a steroid cannot be degraded further (11). Therefore, the formation of these hydroxylated steroids could be assumed to be side reactions, too. However, the fact that most of these compounds could be further degraded by the strain Chol11 wt contradicts this assumption. Even more, the formation of CoA-esters of two of these steroids (P3 and P4) by the acyl-CoA ligase SclA, which is specific for the Δ^4,6^-variant of bile-salt degradation (12), further supports the physiological relevance of these hydroxylated compounds.

For the dihydroxylated compound (P2) the position of one hydroxy group could certainly be assigned to the C9-position of the steroid skeleton because of its UV-spectrum that is characteristic for 9,10-*seco* steroids (11). In contrast, compounds P3 and P4 had UV spectra specific for steroids with a Δ^1,4^-3-keto structure of the steroid skeleton and one hydroxy group. Comparison with authentic standards excluded that these hydroxy groups were located at carbon atoms 3, 6, 7, 9 or 12 of the steroid skeleton. Given that the hydroxylation is part of the metabolic pathway for bile-salt degradation, a monooxygenase must be involved. As the Rieske monooxygenase Nov2c228 is specifically upregulated in cells grown with cholate and deoxycholate compared to growth with 12β-DHADD without side chain (13 (preprint)), a specific role for side-chain degradation is strongly suggested. In support of this hypothesis, Nov2c228, which is quite different from the apparent 9α-hydroxylating Rieske monooxygenases of strain Chol11, produced two metabolites that had an additional hydroxy group at an unknown position when HOCDA was supplied to the *kshA* deletion mutant of *P. stutzeri* Chol1.

The formation of DHSATD by the three Rieske monooxygenases Nov2c407, Nov2c430, and Nov2c440 plus the formation of the apparent *seco*-steroid P2 strongly suggest that bile-salt degradation via the Δ^4,6^-variant also proceeds via the 9,10-seco pathway. Although this was assumed before, 9,10-seco-intermediates had not been detected in this pathway before.

As a prerequisite for the heterologous testing, we identified the KshAB_Chol1_ as a 9α-hydroxylase in Gram-negative bacteria. In contrast to previously identified KshAB monooxygenase systems from Actinobacteria (33, 34), KshAB_Chol1_ only hydroxylated steroids without side chain, which is in agreement with mutants defective in side-chain degradation not producing any *seco*-steroids (16, 17, 20, 22). Additionally, KshAB_Chol1_ was shown to be involved in conversion of HATD (X in Fig 1) to both DHSATD (XI) and THADD (XXIII in Fig S13). Thus, KshAB_Chol1_ performs the formerly known side reaction yielding THADD (11) besides its native function of 9α-hydroxylation. It remains unclear, if both hydroxylations necessary for THADD production are catalyzed by KshAB alone. Other Rieske monooxygenases have been shown to catalyze different reactions with the same substrate depending on binding position and also several subsequent hydroxylations on one substrate molecule upon repeated binding (35–37).

Although the 9α-hydroxylases from strain Chol11 could not be studied in the same detail as KshAB_Chol1_, differences between these proteobacterial enzymes were evident. First, 9α-hydroxylase activity of KshA homologs from strain Chol11 was exclusively detected with *P. stutzeri* Chol1 Δ*kshA* as a host and HOCDA (V) as substrate indicating that a Δ^4,6^- structure was a prerequisite for this reaction. For KshAB_Chol1_ in contrast, both Δ^4,6^- and Δ^1,4^-structures could serve as substrates. Second, the formation of the *seco*-steroid P2, which had an C_5_-side chain, indicates that one of these enzymes presumably catalyzed the 9α-hydroxylation of steroids with side chain; this activity was not observed with KshAB_Chol1_. Third, the 9α- hydroxylases from strain Chol11 as well as the fourth Rieske monooxygenase Nov2c288 did not appear to have specific reductase subunits as KshA_Chol1_ has with KshB_Chol1_. The only putative reductase component from strain Chol11, Novbp123, did not stimulate the activity of any of the KshA homologs from both *P. stutzeri* Chol1 and strain Chol11 in *E. coli*. Novbp123 is expressed constitutively and encoded in a cluster of membrane and electron transport proteins (13 (preprint)). This might indicate a different electron shuttling mechanism for KshA and further enzymes in strain Chol11. The slight activities of Nov2c407, Nov2c430, and Nov2c440 in the *P. stutzeri* Chol1 background therefore must rely on another (unspecific) reductase, which is not present in *E. coli* MG1655.

The apparent dependence of key enzymes for steroid degradation in strain Chol11 on a Δ^4,6^-substructure is in agreement with the constricted growth phenotype of the *hsh2*-deletion mutant of strain Chol11 (28). However, P3 and P4 detected in this study did not have a Δ^6^-double bond anymore according to their UV-spectra. It might therefore be possible that after elimination of the 7-hydroxy group the double bond is reduced similar to the reductive dehydroxylation in *Clostridium* strains (5, 38) (Fig 4D). All three homologs (Nov2c19=5β-Δ^4^-KSTD1, Nov2c85, and Nov2c314) of BaiH, which catalyzes this reaction in *C. scindens* (38), had no activity towards HOCDA with NADH as electron donor when expressed in *E. coli* (data not shown). It is unclear if this reaction is a side-reaction, that does not hinder further degradation as these compounds are degraded by strain Chol11 wt, or a part of regular bile-salt degradation. As HATD (X in Fig 1) can be found in supernatants of strain Chol11 cultures, the B-ring should not be reduced prior to side-chain degradation during unhindered degradation. The lack of homologs for steroid hydratases, aldolases, and thiolases as well as of a second ACAD for C_3_-side chain degradation raises the question whether strain Chol11 cleaves off the whole C_5_-side chain by a so-far unknown mechanism that might involve a monooxygenase reaction catalyzed by Nov2c228 (Fig 10). A hydroxylation at C17 is required at least at one point during cholate degradation in strain Chol11 because HATD (X in Fig 1) is an intermediate of this pathway. Interestingly, the dead-end metabolites produced by *P. stutzeri* Chol1 Δ*kshA* expressing *nov2c228* with HOCDA as substrate had C_3_-side chains. Also, the additional metabolites HOPTC (XXI in Fig 9) and P11 accumulated that could normally be further degraded by *P. stutzeri* Chol1. This suggests that the additional hydroxylation had an inhibiting effect on the second step of side-chain degradation in *P. stutzeri* Chol1. Although this remains a speculation at the current stage, a hydroxy group at C17 might cause such inhibition because of its vicinity to the side chain. On the other hand, a hydroxy group at this position is also necessary for side-chain cleavage, and is inserted by Shy2 followed by aldolytic cleavage by Sal2 in strain Chol1 (20). Although this reaction sequence has been studied in detail in actinobacteria the stereochemistry of the C17 hydroxy group is not known and could only be predicted so far (39–41). Therefore, an unsuited orientation of a hydroxy group at this position may prevent the aldolytic cleavage by Sal2 in strain Chol1. For elucidating side-chain cleavage, a deletion mutant of Nov2c228 would be very useful but, despite numerous attempts, we did not succeed so far with this specific gene. The only enzymes within the side-chain degradation cluster that could have further functions in side-chain degradation are the amidase Nov2c229 and the putative hydroxysteroid dehydrogenases Nov2c226 and Nov2c231. Of these, only Nov2c229 is conserved in the putative side-chain degradation gene clusters of other bile-salt degrading Sphingomonads (*Sphingobium herbicidovorans* MH, *Novosphingobium tardaugens* NBRC16725, and *Novosphingobium aromaticivorans* F199) (13 (preprint)) and therefore is an interesting candidate in our ongoing studies.

**Fig 10.**
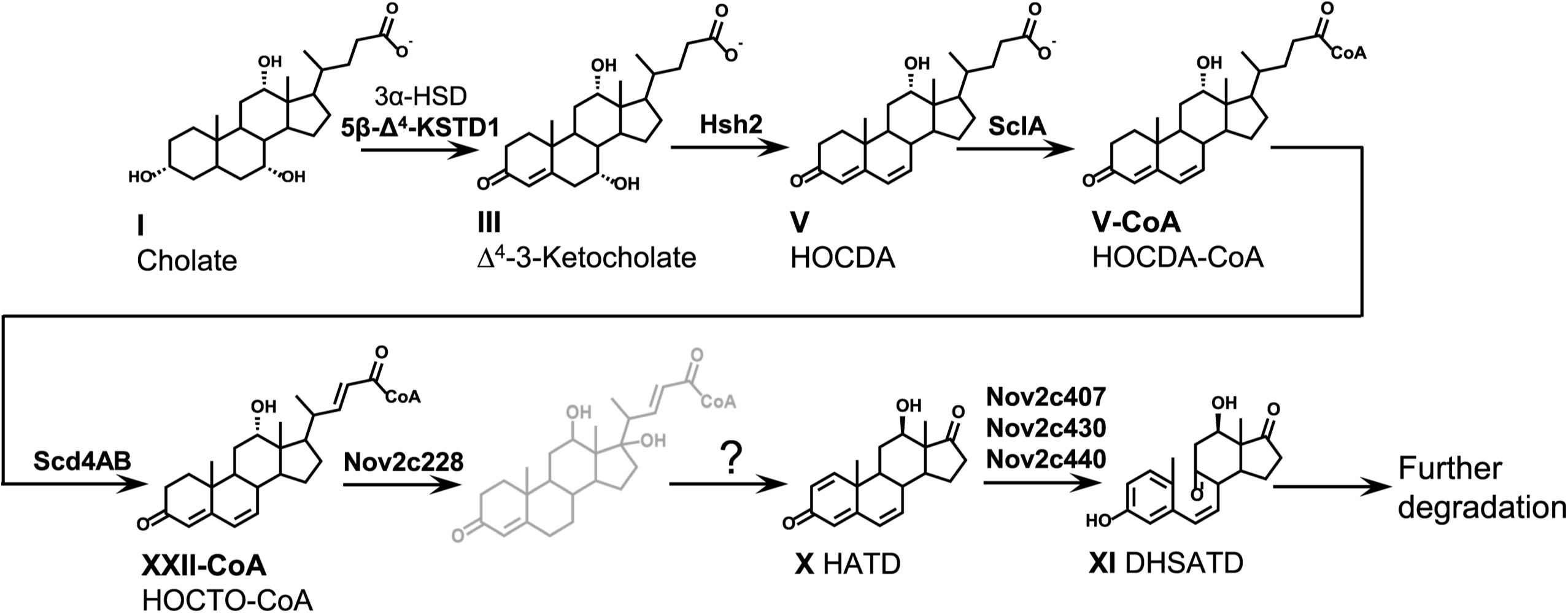
Proposed pathway for A-ring oxidation, B-ring cleavage, and side-chain degradation during cholate degradation in *Sphingobium* sp. strain Chol11. Grey: Structure suggestion. Bold: known enzymes.

## Material and Methods

### Cultivation of bacteria

Strains of *Sphingobium* sp. strain Chol11 (DSM 110934) (11), *P. stutzeri* Chol1 (DSM 103613) (23), and *E. coli* MG1655 (DSM 18039) (42) were grown in the HEPES buffered mineral medium MB as described previously (11, 43). *E. coli* ST18 (DSM 22074) (44) was grown in lysogeny broth medium (LB) (45) with 50 μg ml^−1^ 5- aminolevulinic acid. For strain maintenance and if not indicated otherwise, *P. stutzeri* Chol1 and strain Chol11 wt were grown with 1 mM cholate (I in Fig 1) as carbon source, mutants of *P. stutzeri* Chol1 were grown with 12 mM succinate and *E. coli* MG1655 and mutants of strain Chol11 were grown with 15 mM glucose. Strains containing pDM4 (46) (30 or 90 μg ml^-1^ chloramphenicol), pEX18AP (47) (100 μg ml^-1^ ampicillin for *E. coli* strains or 100 μg ml^-1^ carbenicillin for *P. stutzeri* Chol1), or pBBR1MCS-5 (48) (20 μg ml^-1^ gentamicin) were maintained on LB agar with respective antibiotics and otherwise cultivated in the aforementioned medium with the respective antibiotics. During growth experiments and cultivation with steroids, antibiotics were omitted. Strains were maintained on agar plates prepared from the aforementioned media with 1.5% (w/v) Bacto agar (BD, Sparks, USA).

Liquid cultures up to 5 ml were incubated in 10 ml test tubes and at 200 rpm, whereas larger cultures were incubated in 500 ml Erlenmeyer flasks without baffles. Except for strain maintenance of *E. coli* strains at 37 °C, all strains were cultivated at 30 °C.

Growth experiments were performed in 3 – 5 ml medium in 10 ml test tubes at 30 °C and orbital shaking (Minitron or Ecotron, Infors HT, Einsbach, Germany). Starter cultures for growth experiments of *P. stutzeri* Chol1 or strain Chol11 were grown with succinate for 15 h or glucose for 20 h, respectively, and with antibiotics where appropriate. Main cultures were inoculated from starter cultures without previous washing at an OD_600_ of about 0.02. Growth was monitored by measuring the OD_600_ (Camspec M107, Spectronic Camspec, United Kingdom). At suitable time points, samples for HPLC-MS measurements were withdrawn.

### Biotransformation experiments

Biotransformation of different bile salts by strain Chol11 Δ*scd4A* was determined in 5 ml or 100 ml cultures with 15 mM glucose and 1 mM of the respective bile salt, inoculated from pre-cultures similarly to growth experiments and incubated at 30 °C for two weeks.

For determining biotransformation of the resulting metabolites by strain Chol11, the supernatants of the aforementioned biotransformations were filtered and mixed 1:1 with fresh MB. The resulting medium was inoculated with strain Chol11 from starter cultures and incubated at 30 °C for four days.

For tracking biotransformation of cholate by strain Chol11 cells in dense cell suspensions, 100 ml main cultures with 15 mM glucose were inoculated from starter cultures with glucose and incubated for about 40 h. Cells were harvested by centrifugation (8,000 x g, 4 °C, 8 min), washed with MB without carbon source and resuspended with an OD_600_ of about 1 in MB. 5 ml aliquots of cell suspensions were prepared in 10 ml-reaction tubes and 1 mM cholate was added. Samples for HPLC-MS measurements were withdrawn directly after addition of cholate and at defined time points thereafter.

For testing transformation of HOCDA (V in Fig 1) by *P. stutzeri* Chol1 Δ*kshA* strains and transformation of various steroid compounds by *E. coli* MG1655 strains, cell suspensions with an OD_600_ of about 1 were prepared from starter cultures by washing with and resuspending in MB. 12 mM up to 24 mM succinate were added to suspensions of *P. stutzeri* Chol1 strains and 15 mM up to 30 mM glucose were added to *E. coli* suspensions. The suspensions were incubated for two to four days for *E. coli* MG1655 or six days for *P. stutzeri* Chol1 at 30 °C and 200 rpm orbital shaking. After incubation, cultures were directly frozen at -20 °C and thawed for HPLC-MS measurements.

The ability of SclA to CoA-activate the metabolites P1-P4 produced by strain Chol11 Δ*scd4A* was tested in enzyme assays with cell extracts of *E. coli* MG1655 pBBR1MCS-5::*sclA* (12) or an empty vector control *E. coli* MG1655 pBBR1MCS-5. Cell extracts were prepared from 50 ml – 100 ml cultures in LB with gentamicin by sonication after washing and resupending in 50 mM MOPS buffer (pH 7.8 with NaOH) as described (27) and stored at -20 °C. The protein concentration in the cell free extracts was determined with a BCA assay kit (Pierce, Thermo Scientific, Rockford, IL, United States). Enzyme assays were prepared as follows: cell free extract (12 mg ml^-1^ total protein), 0.5 ml cell-free supernatant containing P1-P4, 3 mM MgCl_2_, 2 mM ATP, 2 mM Li_3_CoA, and 50 mM MOPS buffer (pH 7.8 with NaOH, *ad* 1 ml). For controls, ATP, CoA, or cell extract were omitted. Enzyme assays were incubated at 30 °C and samples for HPLC-MS measurements were withdrawn as indicated.

For elucidating the position of the first additional hydroxy group in P3 and P4, P1-P4 were purified P1-P4 containing cultures by solid phase extraction with reversed phase C_18_ columns (Chromabond, Macherey-Nagel, Düren, Germany). Supernatants were acidified to pH 3 with HCl, columns were washed with methanol and equilibrated with H_2_O_MQ_. After loading the compounds on the columns, columns were washed with first H_2_O_MQ_ and additionally with 10 % methanol. The compounds were eluted with methanol, which was evaporated, and solved in H_2_O with NaOH to gain a neutral pH. Purity and approximate concentration was determined by HPLC-MS. Enzyme assays with purified Hsh2 (28) were prepared in 50 mM MOPS (pH 7.8 with NaOH) and with approximately 2 mM steroid compounds in total. Enzyme assays were incubated at 30 °C for up to 20 h and transformation was monitored by HPLC-MS.

### Cloning techniques and construction of deletion mutants

Cloning was performed according to standard procedures and as described elsewhere (27).

Unmarked deletion mutants of strain Chol11 and *P. stutzeri* Chol1 were constructed as described elsewhere (17, 27) using the suicide vectors pDM4 (46) for strain Chol11 and pEX18AP (47) for *P. stutzeri* Chol1. Up- and downstream regions of the respective gene were amplified with primer pairs upfor/uprev and dnfor/dnrev (Table 4), respectively, and assembled by splicing by overlapping extension (SOE) PCR (49) using primer pair upfor/dnrev. Ligation into pDM4 or pEX18AP was checked using the respective primer pair MCS_for/MCS_rev or M13 primers, respectively. After transfer of the resulting plasmid into strain Chol11 or *P. stutzeri* Chol1 by conjugation, the respective primer pair backbone_for/ backbone_rev was used to verify insertion into the genome. Second recombination was forced by cultivation on LB containing 10 % sucrose. Colonies were checked for gene deletion using primer pair upfor/dnrev. After isolation of a pure culture, gene deletion was verified by PCR of genomic DNA and sequencing of the resulting fragment.

**Table 4.**
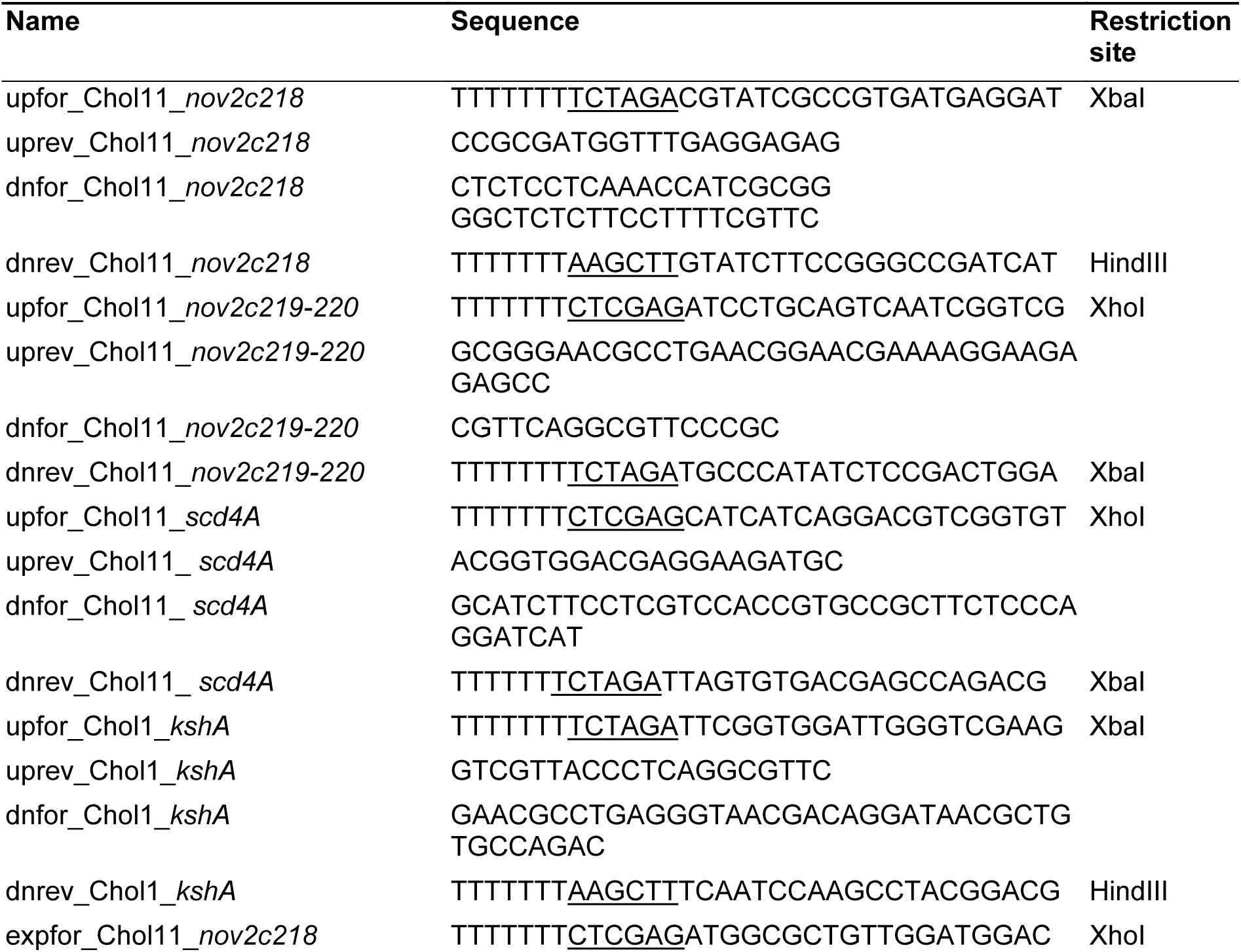
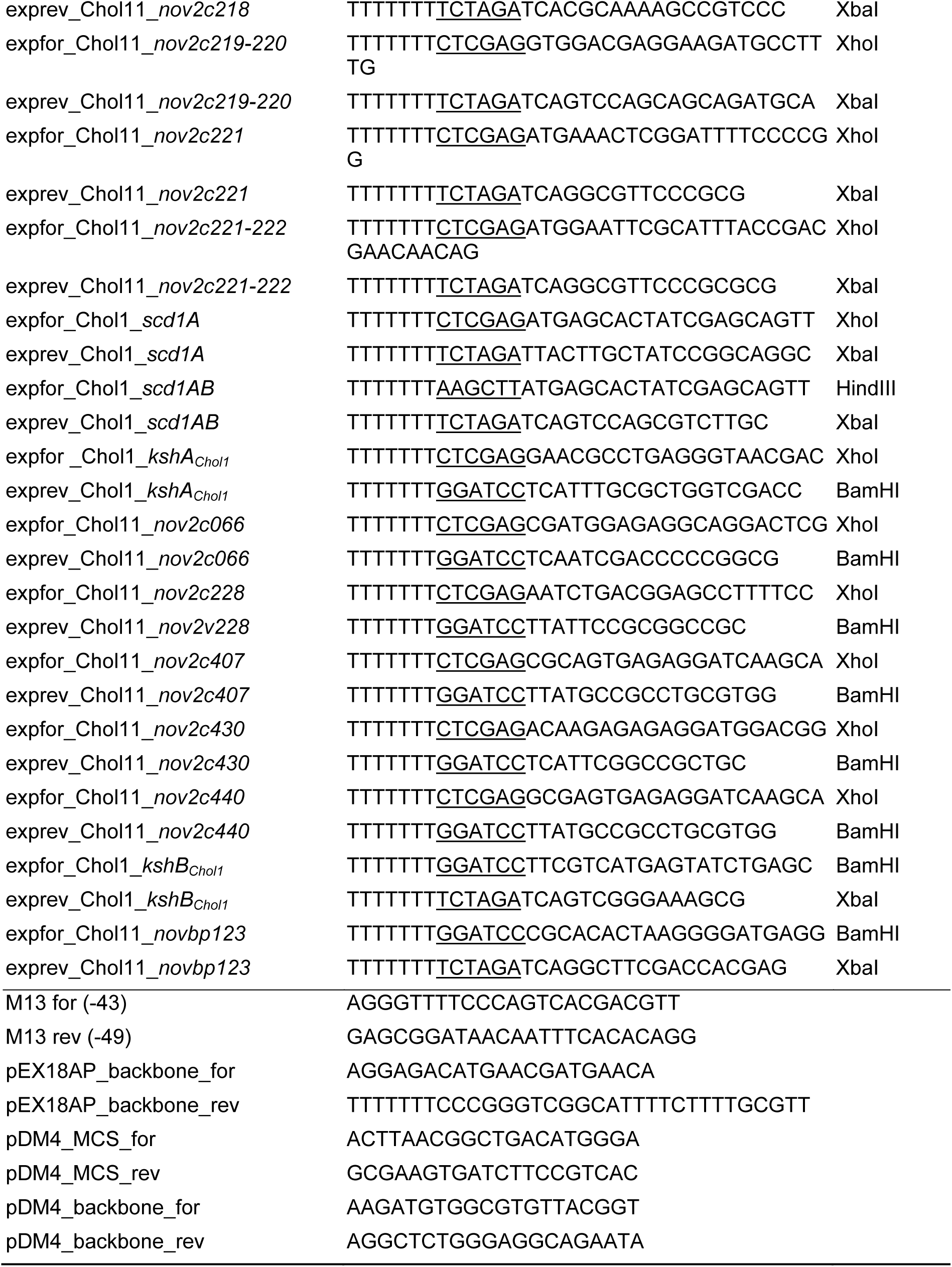
Primers used for construction of unmarked deletion mutants and plasmids for heterologous expression. Underlined: restriction sites

For expression of various genes in *P. stutzeri* Chol1, *E. coli* MG1655, and strain Chol11 strains, genes were amplified using the respective primer pairs expfor/exprev (Table 4) and ligated into vector pBBR1MCS-5. The respective plasmids as well as pBBR1MCS-5::*sclA* (12) were transferred to *E. coli* MG1655 or ST18 by heat shock transformation. For addition of a second gene to the expression vector, the plasmid was isolated and the second gene was added by ligation after restriction. From *E. coli* ST18, vectors were transferred to strains of *E. coli* MG1655, *P. stutzeri* Chol1 and strain Chol11 by conjugation as described (17, 28). Presence of plasmids was confirmed by colony PCR using M13 primers.

HOCDA production strain *P. stutzeri* Chol1 Δ*stdA1* Δ*kstD1* pBBR1MCS-5::*hsh2* was generated by transferring plasmid pBBR1MCS-5::*hsh2* (28) into *P. stutzeri* Chol1 Δ*stdA1* Δ*kstD1* (27).

### Preparation of steroid compounds

Cholate (≥99 %, from ox or sheep bile), deoxycholate (≥97 %), and lithocholate (≥95 %) were purchased from Sigma-Aldrich (St. Louis, MO, USA). Chenodeoxycholate (≥98 %) was purchased from Carl Roth (Karlsruhe, Germany). When lithocholate was added to cultures as a substrate, it was added to MB together with 1 % (w/v) methyl- β-cyclodextrin (TCI, Tokyo, Japan) before autoclaving for solubilization.

Steroid compounds 12β-DHADD (VI), DHOPDC (XII), DHOCTO (XIII) as well as the Δ^4^-3-keto derivatives of bile salts were prepared by biotransformation as described elsewhere (17, 23, 27). All ADDs (VI, XVII, XVIII or XIX) were produced similar to 12β-DHADD by anoxic transformation of the respective bile salt with *P. stutzeri* Chol1. HATD (X) was produced similarly using cholate and *P. stutzeri* Chol1 pBBR1MCS-5::*hsh2* (28). For production of HOCDA (V), *P. stutzeri* Chol1 Δ*stdA1* Δ*kstD1* pBBR1MCS-5::*hsh2* was incubated with cholate and succinate until cholate was completely transformed into HOCDA. For the production of Δ^4^-3-keto bile salts, chenodeoxycholate, ursodeoxycholate or hyodeoxycholate were supplied to *P. stutzeri* Chol1 Δ*stdA1* Δ*kstD1* (27). Dead-end products produced by strain Chol11 Δ*scd4A* from all aforementioned bile salts were produced by biotransformation in 100 ml cultures containing 2 mM of the respective bile salt and incubated for 2 weeks. Δ^6^- HOCTO (XXII) was produced using DHOCTO as a substrate and *P. stutzeri* Chol1 Δ*stdA1* Δ*kstD1* pBBR1MCS-5::*hsh2* for transformation. HOPTC (XXI) was produced like DHOPDC but using HOCDA as substrate.

ADDs, HATD, DHOPDC, DHOCTO, HOCDA, and the Δ^4^-3-keto bile salts were purified by organic extraction as described with dichloromethane for ADDs and HATD or ethyl acetate for the other compounds (21, 23) and resolved in MilliQ pure water (Merck, Darmstadt, Germany). For Δ^6^-HOCTO and the metabolites P1-P8, supernatants of production cultures were sterilized by filtration and used directly in a dilution of 1:1 with fresh medium if not indicated otherwise.

Purity of all steroid compounds was assessed by HPLC-MS measurements. Concentration of HOCDA, DHOPDC, and ADDs was determined photometrically as described (11, 23).

### Analytical methods

Steroid compounds were analyzed by HPLC-MS. Samples were centrifuged (>16,000 x g, ambient temperature, 5 min) to remove cells and particles directly prior to measurement. HPLC-MS measurements were performed using a Dionex Ultimate 3000 HPLC (ThermoFisher Scientific, Waltham, MA, USA) with an UV/visible light diode array detector, coupled to an ion trap Amazon speed mass spectrometer (Bruker, Bremen, Germany) with an electrospray ion source and equipped with a reversed phase C18 column (150 x 3 mm, Eurosphere II, 100-5 C18; Knauer Wissenschaftliche Geräte, Berlin, Germany). For separation, a gradient from 90% to 10 % 10 mM ammonium acetate buffer with 0.1 % formic acid and acetonitrile as described by (27) was used. For analyzing samples of CoA-activation enzyme assays, no formic acid was added to the buffer and measurements were performed at neutral pH.

Cholate concentrations were determined as peak area from base peak chromatograms measured in negative mode. Intermediates were identified due to retention time, UV-, and MS-spectra and comparison with known compounds.

Structures of unknown metabolites were proposed based on retention time as well as UV- and MS-spectra.

### Bioinformatical methods

Homology searches and similarity determinations were performed using the BLASTp algorithm (50, 51). Primer-BLAST (52) was used for generation of primers for construction of deletion mutants. Interpro (53) was used for prediction of protein domains. Alignments and phylogenetic trees were calculated using MegaX (54) (alignment: ClustalW with standard parameters (55), phylogeny: neighbor joining method with standard parameters and 100 bootstrap repetitions (56)) and visualized with iTol (57).

## Acknowledgements

The authors thank Karin Niermann and Kirsten Heuer for excellent experimental support. This work was funded by two grants of the Deutsche Forschungsgemeinschaft (DFG projects PH71/3-2 and INST 211/646-1 FUGG) to BP.

